# Ultrasound neuromodulation reveals distinct roles of the dorsal anterior cingulate cortex and anterior insula in learning

**DOI:** 10.1101/2025.06.12.659273

**Authors:** Nomiki Koutsoumpari, Johannes Algermissen, Siti Yaakub, Hanneke E. M. den Ouden, Nadege Bault, Elsa Fouragnan

## Abstract

Pavlovian biases reflect the notorious influence of hard-wired, evolutionarily conserved cue-response tendencies on instrumental action selection: people show automatic action invigoration in face of potential rewards, but action suppression in face of potential punishments. The neural origin of these biases is unclear. Past evidence suggests dorsal anterior cingulate cortex (dACC) and anterior insula (aIns) as part of a “reset network” that rapidly responds to salient information and might contribute to these biases. We used transcranial ultrasonic stimulation (TUS) in 29 healthy participants to interfere with neural activity in these regions and test their causal role in a within-subject, counter-balanced design across three sessions (sham, TUS-dACC, TUS-aIns). Computational modelling revealed a double dissociation, with distinct roles of both regions in Pavlovian biases: while TUS to the aIns decreased people’s tendency to overly take credit for rewards following action and to ignore punishments following inaction, TUS to dACC increased participants’ tendency to take the cue valence as a reinforcer signal. Although the dACC and aIns are part of the same network and often co-activate during decision-making tasks, TUS interference reveals their distinct roles: the dACC mediates cue-dependence persistence while the aIns is critical for inferring whether outcomes are self-caused.

## Introduction

Some of our everyday decisions are remarkably fast: in the blink of an eye, we paralyze before our hand reaches a potentially venomous spider, or we instinctively reach for the last remaining cookie before our siblings can grab it. Psychology and neuroscience have explained our ability for such fast, instinctive decisions by postulating the existence of multiple decision-making systems that trade-off speed against deliberation and sophistication (Daw et al., 2005; Kahneman, 2011; Keramati et al., 2011). One class of especially fast, but also rigid and seemingly hardwired decisions has been proposed as being driven by a Pavlovian control system (Algermissen and den Ouden, 2023; Dayan et al., 2006; Guitart-Masip et al., 2014; O’Doherty et al., 2017). This system rigidly responds with invigoration to any chance to gain rewards (“Go”), but with inhibition to any looming threat of punishment (“NoGo”). This bias has classically been called the Pavlovian *response bias* (Guitart-Masip et al., 2012; Swart et al., 2017).

The Pavlovian control system gives rise to two other subtle, yet well documented Pavlovian biases that influence learning. One the one hand, a Pavlovian *learning bias* describes people’s tendency to overly claim rewards as caused by their actions, but ignore punishments that follow inaction. For example, a football coach might credit himself for his team’s win after having changed the player lineup, but fail to acknowledge how not changing it led to repeated defeats in the past. One the other hand, a Pavlovian cue-valence dependent *persistence bias* describes people’s tendency to interpret cues that signal the chance for rewards/punishments as reinforcer signals in themselves. In case of no (or an ambiguous) feedback, people are likely to persist in situations in which a reward could be obtained, but likely to change their behaviour in situations in which a negative outcome might occur (Algermissen et al., 2024). Pavlovian biases such as these become evident in tasks that orthogonalize action requirement (Go/NoGo) against the valence of potential outcomes (rewards/punishments) (Guitart-Masip et al., 2012; Swart et al., 2017) and appear to be evolutionarily ancient and shared across the animal realm (Breland and Breland, 1961). By shaping how we assign value to actions and learn from outcomes, these biases play a fundamental role in adaptive behaviour, and their dysregulation may underlie core symptoms across a range of psychiatric disorders (Huys et al., 2016; Scholz et al., 2025).

Pavlovian biases have been linked to neural activity across various cortical and subcortical regions (Algermissen et al., 2024, 2022; Guitart-Masip et al., 2012), yet the precise causal contribution of each of these regions remain unclear. Pinpointing when and where in the brain these biases emerge is essential for uncovering their underlying mechanisms. We have recently observed that the mere anticipation of potential punishment elicits a rapid freezing of gaze within 200–300ms of cue onset (Algermissen and den Ouden, 2024a), coinciding with early value-related components in electroencephalography (EEG) recordings (Algermissen et al., 2024, 2022). These findings suggest that value-related information is processed remarkably early, allowing it to exert a fast and automatic influence on action selection. EEG-informed functional magnetic resonance imaging (fMRI) analyses have further revealed that such early neural responses recruit a network involving the dorsal anterior cingulate cortex (dACC) and anterior insula (aIns), which are selectively engaged by salient/negative information and appear to support rapid behavioural adaptation (Fouragnan et al., 2015). In contrast, later feedback processing stages engage the striatum to support more deliberate reinforcement learning. Together, these findings position the dACC and aIns as strong candidates for causal contributors to Pavlovian biases observed in behaviour.

Selective interference with neural processing in these deep cortical regions to test their role in Pavlovian biases is exceedingly challenging with conventional non-invasive brain stimulation techniques which are limited to superficial cortical regions. However, recently, TUS has emerged as a promising non-invasive tool for neuromodulation (Bault et al., 2024; Murphy and Fouragnan, 2024) that can stimulate deep brain regions with unprecedented spatial precision when careful precautions are taken to limit the transmission loss caused by the skull. TUS has been successfully used to change choice behaviour and neural activity in non-human primates (Folloni et al., 2021; Fouragnan et al., 2019) and humans (Yaakub et al., 2024, 2023b). We thus tested the role of dACC and aIns in Pavlovian biases using a single-blinded within-subject counterbalanced 3-session (dACC/aIns/sham) offline TUS experiment. We expected TUS to interfere with neural processing in these regions and thus lead to different expression of Pavlovian biases in behaviour (Guitart-Masip et al., 2012; Swart et al., 2017) since both regions have been implicated in different aspects of feedback-based learning, including biased credit assignment (Algermissen et al., 2024; de Boer et al., 2019; Swart et al., 2017). We thus also tested for effects on several Pavlovian biases, including a *persistence bias* and a *learning bias*, using computational reinforcement learning models designed to isolate latent cognitive processes.

To foreshadow our results, we found that the stimulation of both dACC and aIns altered Pavlovian biases. Using computational modelling, we found a double dissociation in which dACC and aIns affected distinct learning and decision processes. TUS-aIns selectively reduced participants’ tendency to overly attribute rewards to their own actions and their reluctance to attribute punishments to their inactions, captured by a learning bias parameter (Swart et al., 2017). In contrast, TUS-dACC impaired participants’ ability to tell apart the valence of the feedback from the valence of the cue they responded to, captured by a persistence bias parameter (Algermissen et al., 2024). These results indicate that the dACC plays a role in selecting which cues in the environment to use as a reinforcer signal, whereas the aIns is critical for inferring whether outcomes are self-caused. These two mechanisms — reliance on cue valence as a reinforcer signal and assigning credit to one’s own actions —may be altered in psychiatric disorders (Korn et al., 2014; Shahar et al., 2021) potentially contributing to symptoms such as pathological guilt, externalized blame, and compulsive repetition of actions despite unhelpful outcomes. Disentangling their specific neural contributions is therefore essential for advancing both cognitive neuroscience and clinical psychiatry.

This study also demonstrates the transformative potential of TUS for causally probing decision-making processes in deep cortical regions typically beyond the reach of conventional non-invasive techniques such as transcranial magnetic stimulation (TMS). By targeting the dACC and aIns, two regions often grouped within the salience network and previously implicated in rapid valence-action processing, we show that they play dissociable roles in biased learning. The ability to modulate them independently with TUS represents not only a significant methodological advance but also lays the foundation for mechanistically precise interventions targeting maladaptive decision-making in psychiatric disorders.

## Results

### Participants and procedure

29 participants, screened for any counter indications to TUS and magnetic resonance imaging (MRI) [Contraindications section in Supplementary Material] completed a four-session study protocol. The first session involved collecting MR images for neuronavigation and generating a pseudo-computed tomography (pseudo-CT) image for personalised acoustic planning (Yaakub et al., 2023a) as well as a task practice (Fig. 1A). Participants were then assigned to one of three TUS conditions: Sham, dACC, or aIns (Fig. 1B), using a counterbalanced design. Each of these subsequent sessions included an 80 sec long 5 Hz-patterned TUS intervention, with a spatial peak pulse average intensity (ISPPA) of approximately 50 W/cm^2^ in water, immediately followed by a motivational Go/NoGo learning task (Algermissen et al., 2024; Guitart-Masip et al., 2012; Swart et al., 2017). While the task structure remained consistent across sessions, the cue sets were different and individually randomized to prevent learning effects. Each session lasted approximately 1.5 hours.

**Figure 1.**
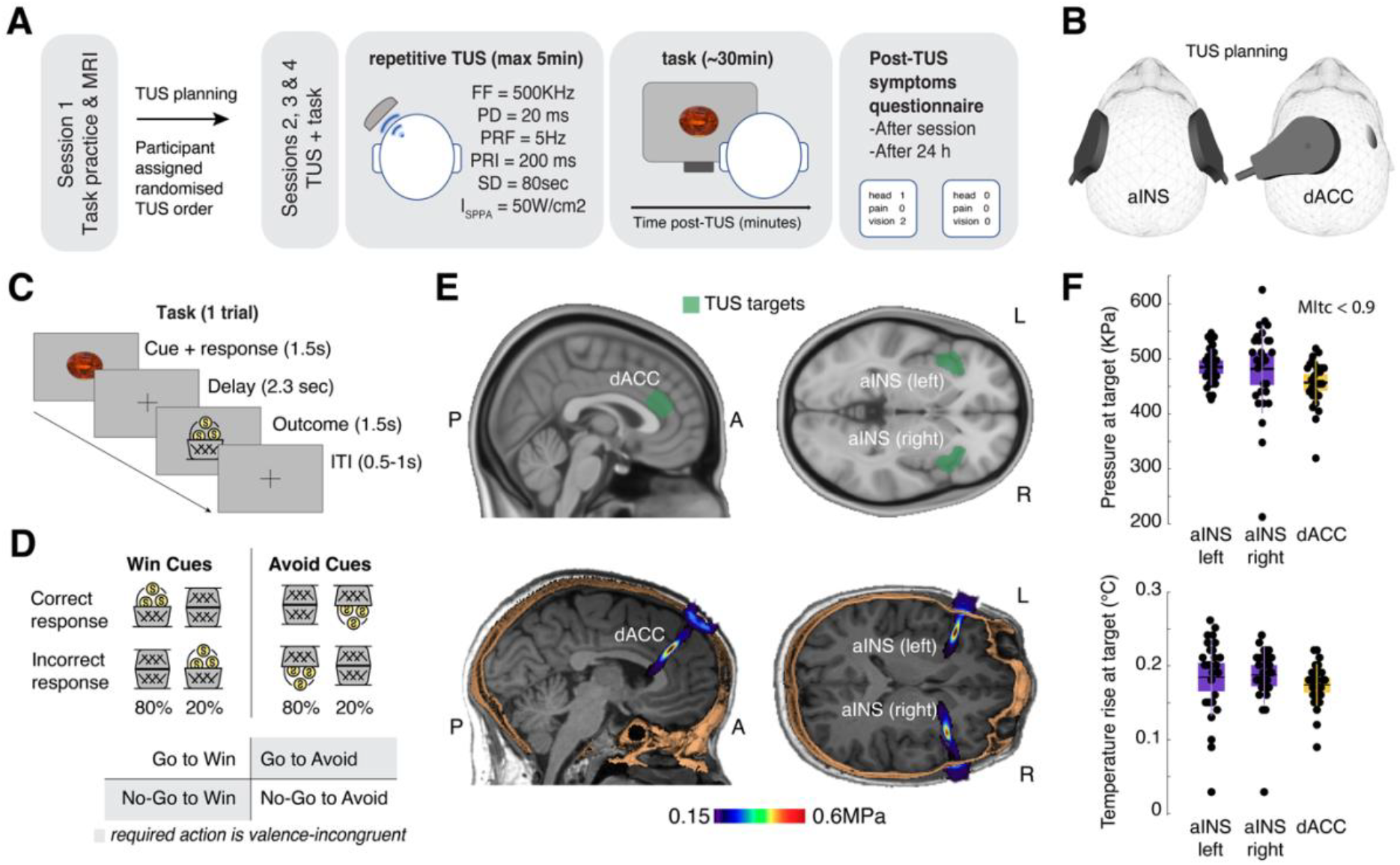
**A**. Study Design: In the first session, a T1-weighted image was acquired for neuronavigation and acoustic planning. Participants also practised a short version of the task. In sessions two to four, participants were randomly allocated to one of three conditions: Sham, dACC, or bilateral aIns. Post-stimulation safety questionnaires were completed on the same day and the following day. **B. TUS positioning:** Visualisation of the positioning of the transducer on the head for the active TUS sessions. **C. Task design:** On each trial, participants saw an abstract cue and needed to learn from trial-and-error to respond with either a Go or NoGo response to that cue. The cue disappeared at the response onset, followed by a jittered interval and finally response-dependent feedback. Each cue had one correct action (Go or NoGo) and was either a Win cue (leading to rewards or neutral outcomes) or an Avoid cue (leading to neutral outcomes or punishments). Rewards and punishments were represented as money falling into or out of a bucket. **D. Feedback validity and cue types:** Each block of session included 4 novel cues, varying in valence (Win/Avoid) and required action (Go/NoGo). Incongruent cues, where the required action was in opposition to the Pavlovian bias (NoGo to Win, Go to Avoid punishment), are highlighted in grey. Feedback was probabilistic. Correct responses to Win cues resulted in rewards 80% of the time and neutral outcomes 20% of the time. Correct actions to Avoid cues led to neutral outcomes in 80% of cases and punishments in 20%. Incorrect responses led to feedback with the reverse probabilities. **E. Regions:** Targeted brain regions for TUS (dACC and bilateral aIns) are shown (top), with post-stimulation pressure simulated through k-Plan software (BrainBox, Inc.) expressed in megapascals on a representative participant (bottom). **E. Regions:** Targeted brain regions for TUS (dACC and bilateral aIns) are shown (top), with post-stimulation pressure simulated through k-Plan software (BrainBox, Inc.) expressed in megapascals on a representative participant (bottom). **F. TUS simulations:** Maximum pressure in the brain target volume (top panel) and maximum temperature rise in the brain target volume (bottom panel), (n=29). Box plots show the mean, and the standard error (bounds of the box). Data from each individual participant are presented as small black circles.

During the motivational Go/NoGo learning task, on each trial, participants were presented with one of several cues. They had to learn from probabilistic feedback via trial-and-error to either perform a Go (button press) or a NoGo action (no button press) to four cues per block. Half of these cues were Win cues for which correct responses (mostly) returned a reward (increase in point score), while incorrect responses (mostly) led to neutral feedback (no change in point score). For the other half of cues, called Avoid cues, correct responses (mostly) led to neutral feedback, while incorrect responses (mostly) led to punishments (decrease in point score). The task timeline is presented in Fig. 1D. Feedback was probabilistic such that correct responses resulted in the desired outcome (reward or omitted punishment) 80% of the time, whereas incorrect actions led to the desired outcome in only 20% of trials. On each session, participants completed 320 trials, split into four blocks of 80 trials. In each block, a distinct set of four cues was presented, each repeated 20 times. The order of cue presentations was randomized for each participant within each block (Fig. 1D).

Targeting of the bilateral aIns and dACC was individually adjusted using T1-weighted MRI (structural scans) during planning for target and transducer placement, as well as acoustic and thermal simulation. The dACC was defined anatomically as the region anterior to the anterior commissure, dorsal to the genu of the corpus callosum, and bounded dorsally by the cingulate sulcus. The aIns was identified as the anterior portion of the insular cortex, located deep within the Sylvian fissure, anterior to the central insular sulcus, and bordering the frontal operculum. Repetitive TUS followed a 5 Hz-patterned protocol with a 10% duty cycle (Fundamental Frequency = 500 KHz, Pulse Duration = 20 ms, pulsed every 200 ms), applied for 80 seconds (400 pulses in total). The ISPPA in water was maintained at 50 W/cm^2^ across participants (validated in a hydrophone tank (Yaakub et al., 2023b)). We took care to remain within guidelines for human ultrasound exposure as defined the International Thermal and Radiological Ultrasound Safety Standards and Thresholds (ITRUSST; Aubry et al., 2024). Importantly the transcranial Mechanical Index (MItc) was way below 1.9 and the results of our acoustic simulations predominantly indicated a maximum soft tissue temperature rise below 2°C [Supplementary Table 1]. We also calculated the Cumulative Equivalent Minutes at 43°C (CEM43), a metric reflecting both duration and intensity of heating relative to 43°C, the critical threshold for thermal cell damage. The CEM43 values were always below 0.1, i.e. well below the safety threshold of 0.25 (Aubry et al., 2024).

Sham was administered in the same manner as active aIns-TUS, placing the transducer over the site but without actual stimulation. Instead, participants listened to a sound that mimicked the TUS protocol’s acoustic output (validated across 6 participants independent from this study), played through bone conduction headphones. Acoustic simulations for each participant were used to plan the stimulation to ensure accurate pressure delivery at the target area and adherence to safety protocols (an example participant’s targeted areas are shown in Fig. 1E). Detailed acoustic simulation parameters and outputs for all study participants can be found in Fig. 1F, Supplementary Materials [Table 1] and summarised in Table 1.

**Table 1.** Ultrasound Parameters Across Different Target Tissues for dACC and aIns Stimulation. A sphere (Target ROI) with a 3mm radius centred on the target coordinates is used to reliably extract peak values from the brain region where the focus is located.

| Target | Tissue | Temperature (°C) |  | Pressure (MPa) |  | ISPPA (W/cm<sup>2</sup>) |  | MI<sub>tc</sub> |  |
| --- | --- | --- | --- | --- | --- | --- | --- | --- | --- |
|  |  | *M* | *SD* | *M* | *SD* | *M* | *SD* | *M* | *SD* |
| dACC | Target ROI | 37.18 | 0.05 | 0.45 | 0.04 | 6.69 | 1.16 | 0.63 | 0.06 |
|  | Soft tissue | 37.96 | 0.28 | 0.45 | 0.04 | 6.72 | 1.18 | 0.63 | 0.06 |
|  | Skull | 38.73 | 0.41 | 0.32 | 0.08 | 3.57 | 2.29 | 0.45 | 0.12 |
| l-aIns | Target ROI | 37.19 | 0.04 | 0.47 | 0.08 | 7.60 | 2.26 | 0.67 | 0.11 |
|  | Soft tissue | 38.14 | 0.34 | 0.48 | 0.06 | 7.74 | 1.98 | 0.68 | 0.09 |
|  | Skull | 38.99 | 0.64 | 0.33 | 0.05 | 3.63 | 1.09 | 0.46 | 0.07 |
| r-aIns | Target ROI | 37.17 | 0.03 | 0.48 | 0.03 | 7.53 | 1.06 | 0.67 | 0.05 |
|  | Soft tissue | 38.20 | 0.38 | 0.48 | 0.03 | 7.54 | 1.05 | 0.67 | 0.05 |
|  | Skull | 39.10 | 0.62 | 0.35 | 0.06 | 4.06 | 1.39 | 0.49 | 0.08 |

### Side effects

Participants completed a post-TUS questionnaire for side effects twice: once at the end of each session, and a second time 12 to 48 hours after the session. They were presented with a list of symptoms for which they had to rate the intensity of the symptom and the degree to which they believe that symptom was related to the stimulation (refer to the Methods section under TUS protocol and procedure for further safety details; see also Supplementary Material 1 for the full safety questionnaire). Immediately after stimulation of the insula, participants reported the following side effects: anxiety (N = 1), difficulty paying attention (N = 2), forgetfulness (N = 1), headache (N = 1) and sleepiness (N = 3). All side effects were rated as mild and unrelated to the stimulation. Following stimulation of the dACC, moderate anxiety (N = 1), mild forgetfulness (N = 1), moderate neck pain (N = 1), mild to moderate sleepiness (N = 2) and mild vision problem (N = 1) were reported. Only the participant reporting vision problems considered those as possibly related to the stimulation. Following sham stimulation, one participant reported mild itchiness (N = 1) and another moderate sleepiness (N = 1), again both problems were rated as unrelated to the stimulation. Overall, no qualitative difference in the number and quality of side effects was observed between sham and active stimulation. In most cases of side effects, participants did not attribute them to the stimulation.

### Pavlovian biases in responding and learning in the sham session

We first focus on the data from the sham session to replicate Pavlovian biases in responding and learning as typically observed in this task (Algermissen et al., 2024; Guitart-Masip et al., 2012; Swart et al., 2017).

First, in line with previous work (Algermissen et al., 2024; Scholz et al., 2022; Swart et al., 2017; van Nuland et al., 2020), we used mixed-effects logistic regression to fit Go/NoGo responses as a function of the required action of a given cue (Go vs. NoGo), the cue valence (Win vs. Avoid cues), and their interaction. Participants successfully learned the task, as they made more Go responses to Go than NoGo cues (main effect of required action: *b* = 2.132, 95%-CI [1.797, 2.467], χ^2^ (1) = 53.812, *p* <.001; Fig. 2D). Furthermore, there was a Pavlovian bias in behaviour, indexed by significantly more Go responses to Win than Avoid cues (main effect of cue valence: *b* = 0.689, 95%-CI [0.421, 0.966], χ^2^ (1) = 17.173, *p* <.001). Lastly, the interaction between both terms was significant (*b* = 0.269, 95%-CI [0.104, 0.434], χ^2^ (1) = 9.646, *p* =.002), reflecting a stronger Pavlovian bias for Go than for NoGo cues. Similar analyses of the reaction times (RTs) also revealed a bias (Supplementary Fig. 2). These analyses confirm that participants’ behaviour was influenced by Pavlovian biases in the sham session, which constitutes a prerequisite for modulating them in the other two active sonication sessions.

**Figure 2.**
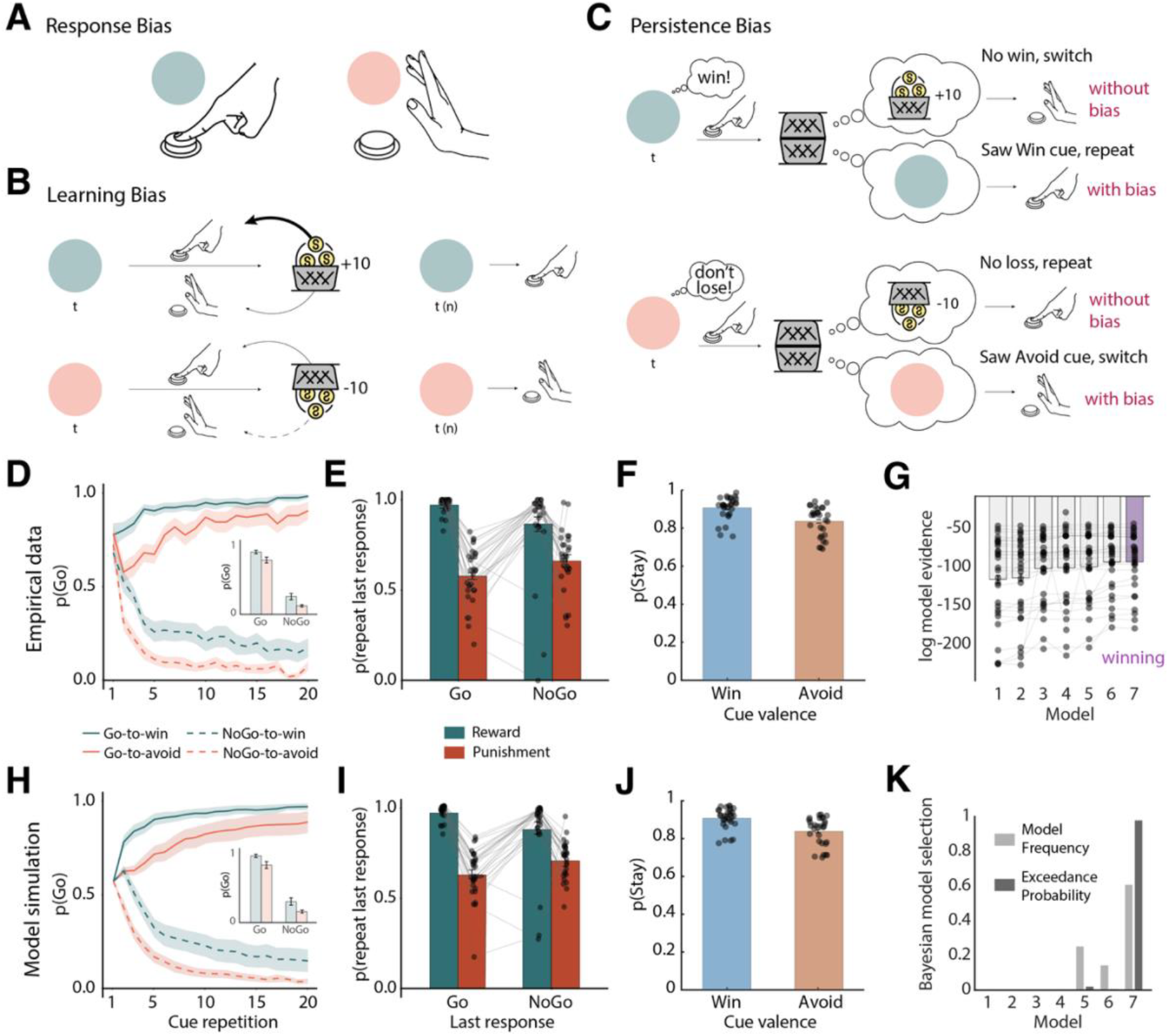
Behavioural performance in the sham dataset. **A. Response bias:** Humans exhibit a tendency to show actions (i.e. button presses) for Win cues (blue), aligning reward seeking with action invigoration, and to withhold actions for Avoid cues (red), aligningpunishment avoidance with action suppression/inaction. **B. Learning bias:** Learning is enhanced for rewarded Go actions compared to rewarded NoGo actions, promoting faster acquisition of Go actions from rewards. Conversely, learning is reduced for punished NoGo actions compared to punished Go actions, slowing the extinction of NoGo responses via punishments. **C. Persistence bias:** Seeing a Win cue partly acts as a reward in itself, leading to a higher tendency to repeat responses to such cues, even when a neutral outcome (negative feedback) suggests that one should switch the response. In contrast, seeing an Avoid cue partly acts as a punishment in itself, leading to a higher tendency to switch responses, even if a neutral outcome (positive feedback) suggests repeating the response. **D**. Trial-by-trial proportion of Go responses (error bands = ± SEM, n = 29) for Go cues (solid lines) and NoGo cues (dashed lines). A clear response bias is evident from the start, with participants making more Go responses to Win (green lines) than Avoid (red lines) cues. Participants show a general bias towards Go responses, with initial response rates around 80%. Additionally, instrumental learning is apparent in Go responses increasing for Go cues and decreasing for NoGo cues over time. **E**. Probability to repeat a response on the next encounter of the same cue as a function of the response and outcome for valenced outcomes (rewards, punishments) only (error bars are ±SEM across participants, n = 29, dots indicate individual participants). Learning was reflected in the higher probability of staying after rewarded responses than after being punished (main effect of outcome valence). Biased learning was evident in the stronger outcome effect after Go responses than after NoGo responses, indicating the presence of a **l**earning bias. **F**. Probability to repeat a response on the next encounter of the same cue as a function of cue valence (error bars are ± SEM across participants, n = 29, dots indicate individual participants). Participants showed a higher tendency to repeat responses for Win than Avoid cues, indicative of a *persistence* bias. **G**. Laplace approximation to the log-model evidence favours the model incorporating all three Pavlovian biases—response bias, learning bias, and persistence bias (M7)—over simpler models (M1–M6); error bars are ± SEM across participants, n = 29. **H-J**. Posterior predictive checks: one-step-ahead predictions based on the best fitting parameters (estimated with hierarchical Bayesian inference) for M7. **K**. Bayesian model selection. The winning model is M7 as determined by model frequency and protected exceedance probability, in line with log model evidence.

Second, we used mixed-effects logistic regression to analyse participants’ propensity to repeat vs. switch a response on the next trial as a function of the obtained outcome and performed response on the current trial. We have previously found that Pavlovian biases are present not only in action selection, but also in learning, with a larger propensity to take valenced feedback into account (i.e., repeat actions after rewards, switch actions after punishments) for Go actions compared to NoGo actions. Specifically, learning is increased for rewarded Go (relative to rewarded NoGo) actions and decreased for punished NoGo (relative to punished Go) actions (see detailed explanation in the computational modelling) (Algermissen et al., 2024; Swart et al., 2017). Because this bias should only occur for valenced outcomes (rewards, punishments), but not for neutral outcomes, we only included valenced outcomes in this analysis. We observed a significant main effect of the obtained outcome (reward vs. punishment), *b* = 1.436, 95%-CI [1.190, 1.683], χ^2^ (1) = 50.170, *p* <.001, with more response repetitions after rewards than after punishments, and a main effect of performed response, *b* = 0.211, 95%-CI [0.081, 0.341], χ^2^ (1) = 6.422, *p* =.001, with more response repetitions for Go than NoGo responses (likely due to the fact that participants showed overall more Go than NoGo responses in this task). Crucially, the interaction between the obtained outcome and performed response was significant, *b* = 0.423, 95%-CI [0.195, 0.650], χ^2^ (1) = 10.051, *p* <.001, with a stronger outcome effect after Go than after NoGo responses (Fig. 2E). This result confirmed the presence of Pavlovian biases also in learning from feedback.

Thirdly, we used mixed-effects logistic regression to analyse participants’ propensity to repeat vs. switch a response on the next trial as a function of the cue valence. We have previously found that participants show a propensity to repeat responses for Win cues than for Avoid cues, suggesting that they also partly interpret the cue valence as a feedback signal (Algermissen et al., 2024). This leads to response repetition (persistence) tendencies that depend on the cue valence, a tendency that was not well captured by previous computational models (Supplementary Fig. 4), but can only be accounted for by a third, *persistence bias*. Indeed, we found a strong main effect of cue valence on response repetitions, *b* = 0.351, 95%-CI [0.262, 0.440], χ^2^ (1) = 60.117, *p* <.001, with more repetitions for Win than Avoid cues (Fig. 2F). This result indicated the presence of a third Pavlovian bias in which the cue valence affected persistence tendencies. Taken together, these results motivated us to further interrogate these biases using computational reinforcement learning models.

### Computational modelling of the sham data

We next fit a series of increasingly complex reinforcement learning models to participants’ responses in the sham sessions. These models follow previous publications on this task (Algermissen et al., 2024; Guitart-Masip et al., 2012; Swart et al., 2017). These models incorporate three previously described forms of Pavlovian biases: (a) a response bias, (b) a learning bias, and (c) a persistence bias.

The first bias, called ***response bias***, is a bonus/malus added to the learned action values (Fig. 2A): For Win cues, a bonus is added to the value of Go actions, and for Avoid cues, a malus is subtracted from the value of Go actions (Guitart-Masip et al., 2012). This bias is present at the onset of the task and constant throughout learning. Since action values and the bias jointly determine which action is selected, and since action values will increasingly separate with learning over the course of the task, the role of this response bias will, in net, decrease over the time (Swart et al., 2017).

The second bias, called ***learning bias***, captures two tendencies well described in the literature: people ascribe positive outcomes to their own actions (which in other contexts can lead to “illusions of control” (Brown and Jenkins, 1968; Wegner, 2002)), but tend to ignore negative outcomes after having remained passive (a form of “omission bias” (Ritov and Baron, 1990; Zeelenberg et al., 2000); Fig. 2B). These tendencies are captured by a boost in the learning rate for rewarded Go actions and a decrement in the learning rate for punished NoGo actions (Swart et al., 2017). The impact of this bias on behaviour only develops over time, where the action values become increasingly distorted relative to unbiased learning.

The third bias, called ***persistence bias***, captures the fact that the cue valence (Win or Avoid cue) can act as a reinforcer (Fig. 2C) (Algermissen et al., 2024): For Win cues, rewards signal positive feedback, and neutral outcomes signal negative feedback. However, the Win cue itself can erroneously be interpreted as a feedback signal and reinforce behaviour irrespective of the eventual outcome. Thus, while a neutral outcome for a Win cue signals negative feedback, which suggests switching behaviour, the cue valence itself might be taken for positive feedback, which usually translates into a response repetition. Vice versa, for Avoid cues, neutral outcomes signal positive feedback and punishments signal negative feedback. The Avoid cue itself can erroneously be interpreted as negative feedback. Thus, while a neutral outcome for an Avoid cue signals positive feedback, which suggests repeating behaviour, the Avoid cue itself can be taken for negative feedback, which usually translates into a response switch. We have previously observed that this bias can dominate repeat/switch behaviour after neutral outcomes, resulting in the paradoxical observation of more response repetition after negative (i.e. no reward) feedback for Win cues than positive (i.e. no punishment) feedback to Avoid cues (Algermissen et al., 2024). Such a bias should be invisible at the beginning of the task and only arise with learning, leading to increasingly distorted learning curves over time.

We tested the presence of these three biases in behaviour using a series of seven reinforcement learning models. The base model (M1) was a simple Q-learning model that learned the values of Go and NoGo actions for each cue. To generate choices, action values were turned into action probabilities via a softmax transform. Action values were subsequently updated based on the obtained outcomes via reward prediction errors, which were scaled by a free learning rate parameter ε and then added to the old action values. In line with previous implementations (Algermissen et al., 2024; Guitart-Masip et al., 2012; Swart et al., 2017), objective outcomes were scaled by a free feedback sensitivity parameter ρ. High values on this parameter lead to learned action values developing further apart and thus more deterministic choices, similar to a high inverse temperature parameter used in alternative models. We extended this model with a constant Go bias parameter b to capture overall tendencies towards Go or NoGo responses (M2), a response bias parameter π that added a bonus/malus to the action value of Go depending on the valence of a cue (M3), and finally, a learning bias parameter κ capturing increased learning rates for rewarded Go responses and decreased learning rates for punished NoGo responses (M4). M5 combined all three parameters and fitted the data better than any of the simpler models, suggesting that both a response bias and learning bias were present in the data. Finally, we extended M5 by adding a persistence parameter φ_INT_ that captured overall tendencies to repeat vs. switch actions (M6), and a persistence bias φ_DIFF_ that captured the higher tendency to repeat actions for Win cues than for Avoid cues (M7).

We compared these seven models using Bayesian model selection (Stephan et al., 2009). In line with our previous work (Algermissen et al., 2024), M7, implementing all three Pavlovian biases, was the best-fitting model according to the log-model evidence (Fig. 2G), the model frequency (M7: 61%; M6: 14%; M5: 25%), and crucially, according to the protected exceedance probability (M7: 98%; M5: 2%; Fig. 2K). We performed posterior-predictive checks by simulating the action probabilities based on the fitted parameters combined with participants’ actions and outcomes (one-step ahead predictions), which captured the empirical data patterns very well (Fig. 2H, I, J). Simpler models failed to capture core features of the empirical data patterns. Model recovery showed that all models were distinguishable from each other, and parameter recovery on M7 showed that all parameters were well identifiable (Supplementary Fig. 6 & 7). We also explored alternative models in which cue valence did not affect persistence directly, but biased the outcomes in the prediction error updating equation. Such models predicted, qualitatively, a very similar pattern as M7, but showed slightly inferior fit (Supplementary Fig. 5). In sum, these results confirmed the presence of all three Pavlovian biases in participants’ behaviour in the sham sessions.

### Sonication effects on Go/NoGo choices

We next analysed the effect of TUS on Pavlovian biases in responding and learning. We again first used regression models based on participants’ Go/NoGo responses and their propensity to repeat/switch responses, and then subsequently contrasted parameters obtained from the best-fitting reinforcement learning model across all sonication conditions.

We used a mixed-effects logistic regression model to test for effects of TUS on (biased) responding, i.e., their Go/NoGo responses. This model included the required action, cue valence, the TUS sonication condition, as well as the block half (first vs. second half of a block) to account for the possibility that TUS might affect learning processes, which might only become visible in the second half of each block. Beyond the main effects of required action and cue valence and their interaction, this model yielded a significant 3-way interaction between required action, cue valence, and sonication condition, χ^2^ (2) = 14.277, *p* <.001, a significant 3-way interaction between cue valence, sonication condition, and block half, χ^2^ (2) = 9.191, *p* =.010, and a significant 4-way interaction between required action, cue valence, sonication condition, and block half, χ^2^ (2) = 21.548, *p* <.001 (Fig. 3A, B, C). See Supplementary Results for results when analysing each cue condition (Go-to-Win, Go-to-Avoid, NoGo-to-Win, NoGo-to-Avoid) separately. To better understand these effects, we followed them up with the regression analyses of response repetitions vs. switches.

**Figure 3.**
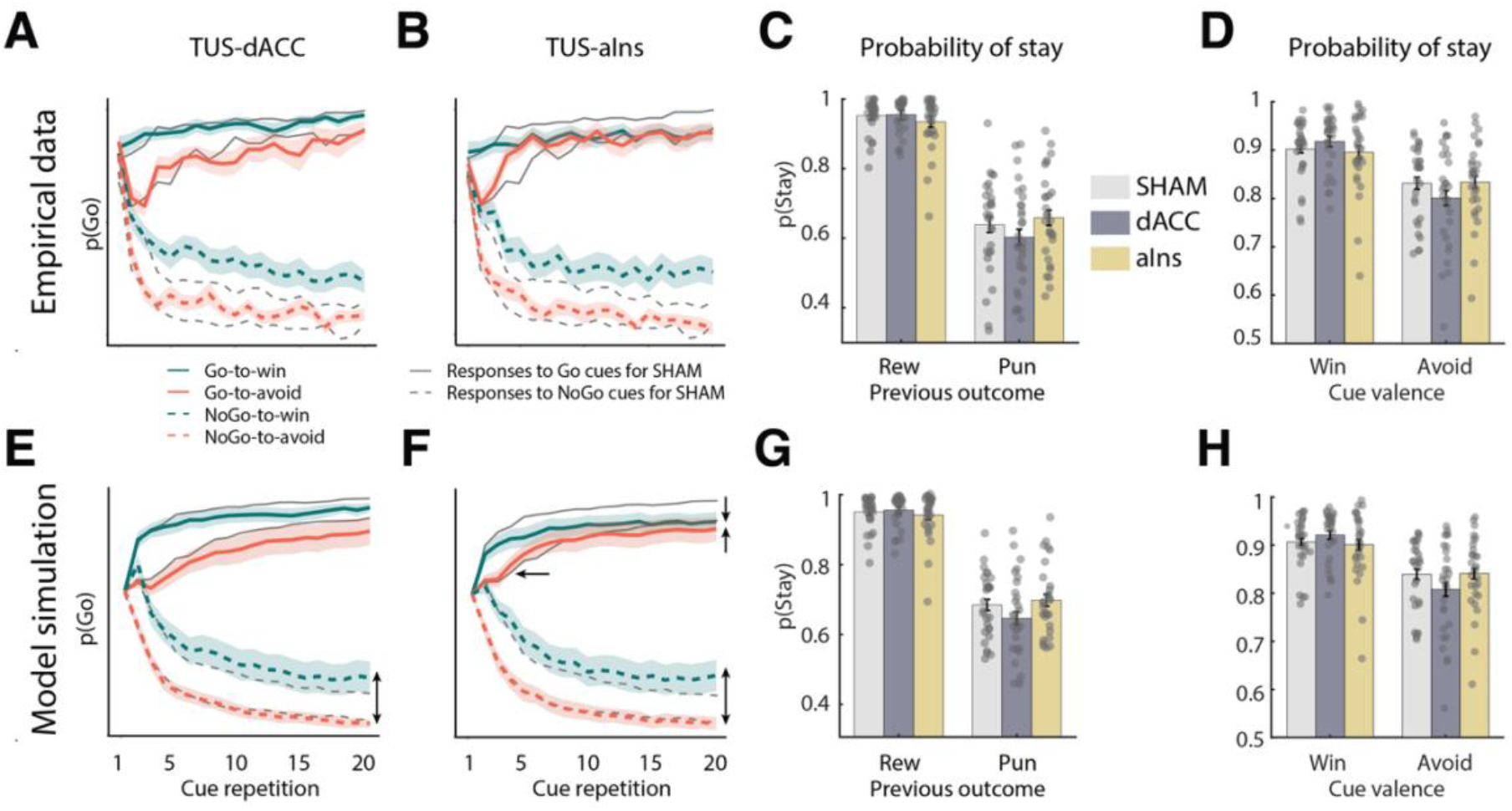
**A-B Empirical Probability of Go**. Trial-by-trial proportion of Go responses across sonication conditions (TUS-dACC, and TUS-aIns, grey lines represent the sham condition). TUS-dACC decreased Go responses in the Go-to-Win condition but increased them in the NoGo-to-Win condition. TUS-aIns decreased Go responses in the Go-to-Win condition but increased them in the Go-to-Avoid and NoGo-to-Win conditions. Coloured lines display the active sonication condition. **C-D Probability of Stay**. TUS effects on response repetitions based on outcomes (rewarded Go vs. punished NoGo) and cue valence (Win vs. Avoid). TUS-aIns (but not TUS-dACC) attenuated the tendency to repeat rewarded Go responses, but increased the tendency to repeat punished NoGo responses compared to sham. TUS-dACC (but not TUS-aIns) increased the cue valence effect on response repetitions, with stronger persistence for Win cues relative to Avoid cues. **E-H** Posterior predictive checks reproduced the empirical data patterns for the probability of Going and staying.

### Sonication effects on response repetitions/switches

Analyses of Go/NoGo choices suggested effects of TUS on learning (i.e., changes in Go/NoGo responses over time). To more directly investigate learning, i.e., the impact of outcomes on subsequent choices, we analysed participants’ repeat/switch choices as a function of outcomes and responses on the current trial. We focused on the signatures of biased learning.

First, we contrasted sonication conditions selectively on those trials on which biased learning should occur, namely trials with rewarded Go responses or with punished NoGo responses. If the learning bias became stronger, participants would repeat rewarded Go responses more and punished NoGo responses less, yielding a stronger difference between both conditions (with no change in any other condition). Vice versa, if the bias became weaker, the difference between both conditions would decrease. In a mixed-effects logistic regression model fit to the response repetitions vs. switches, we observed a significant main effect of outcome (reward vs. punishment) qualified by a significant outcome × sonication interaction, χ^2^ (2) = 6.162, *p* =.046: participants more often repeated rewarded Go responses than punished NoGo responses and this tendency was significantly attenuated after TUS-aIns relative to sham, *b* = -0.150, 95%-CI [-0.293, -0.006], χ^2^ (1) = 4.197, *p* =.040, and not after TUS-dACC compared to sham, *b* = 0.001, 95%-CI [-0.058, 0.249], χ^2^ (1) = 0.001, *p* =.984 (Fig. 3C). These results suggest that TUS-aIns, but not TUS-dACC, reduced the learning bias.

Second, we tested whether sonications changed the effect of cue valence on response repetitions vs. switches, reflecting a bias in persistence. Participants were more likely to repeat responses for Win than Avoid cues (main effect of valence: *b* = 0.428, 95%-CI [0.342, 0.514], χ^2^ (1) = 94.620, *p* <.001], which was qualified by a cue valence × sonication interaction, χ^2^ (2) = 7.617, *p* =.022: the cue valence effect was significantly stronger after TUS-dACC compared 440 to sham, *b* = 0.113, 95%-CI [0.031, 0.194], χ^2^ (1) = 7.368, *p* =.007, with no difference between TUS-aIns and sham, *b* = 0.006, 95%-CI [-0.063, 0.074], χ^2^ (1) = 0.025, *p* =.873 (Fig. 3D). These results suggest that TUS-dACC, but not TUS-aIns, affects the persistence bias. Next, we directly confirmed these observations by fitting these parameters using computational reinforcement learning models and comparing the fitted parameters across TUS sonication conditions.

### Effects of TUS on computational modelling parameters

We fit the best-fitting model from the sham session, M7, comprising all three Pavlovian biases, namely a response bias π, a learning bias κ, and a persistence bias φ_DIFF_, to the other two sonication conditions. Model comparisons suggested that, in the TUS-dACC sessions, M7 was again the best-fitting model according to both the model frequency (M7: 89%; M6: 11%) and the protected exceedance probability (M7: 100%). In the TUS-aIns sessions, a slightly different outcome occurred, with M6 being the best fitting model according to both the model frequency (M6: 68%; M7: 32%) and protected exceedance probability (M6: 97%; M7: 3%). Given that M6 is a nested version of M7 (i.e., M7 with the φ_DIFF_ parameter fixed to zero is equivalent to M6), we proceeded with comparing the parameter values in M7 across all three sonication conditions. We focused on the three parameters reflecting the three different bias parameters: the response bias π, the learning bias κ (Fig. 4A), and the persistence bias φ_DIFF_ (Fig. 4D). We compared the parameter values between all three conditions with a one-way repeated-measures ANOVA, following up any significant effects with paired *t*-tests. There was no significant difference in the response bias π, *F*(2, 56) = 1.719, *p* =.189, generalized η^2^ = 0.018.

**Figure 4.**
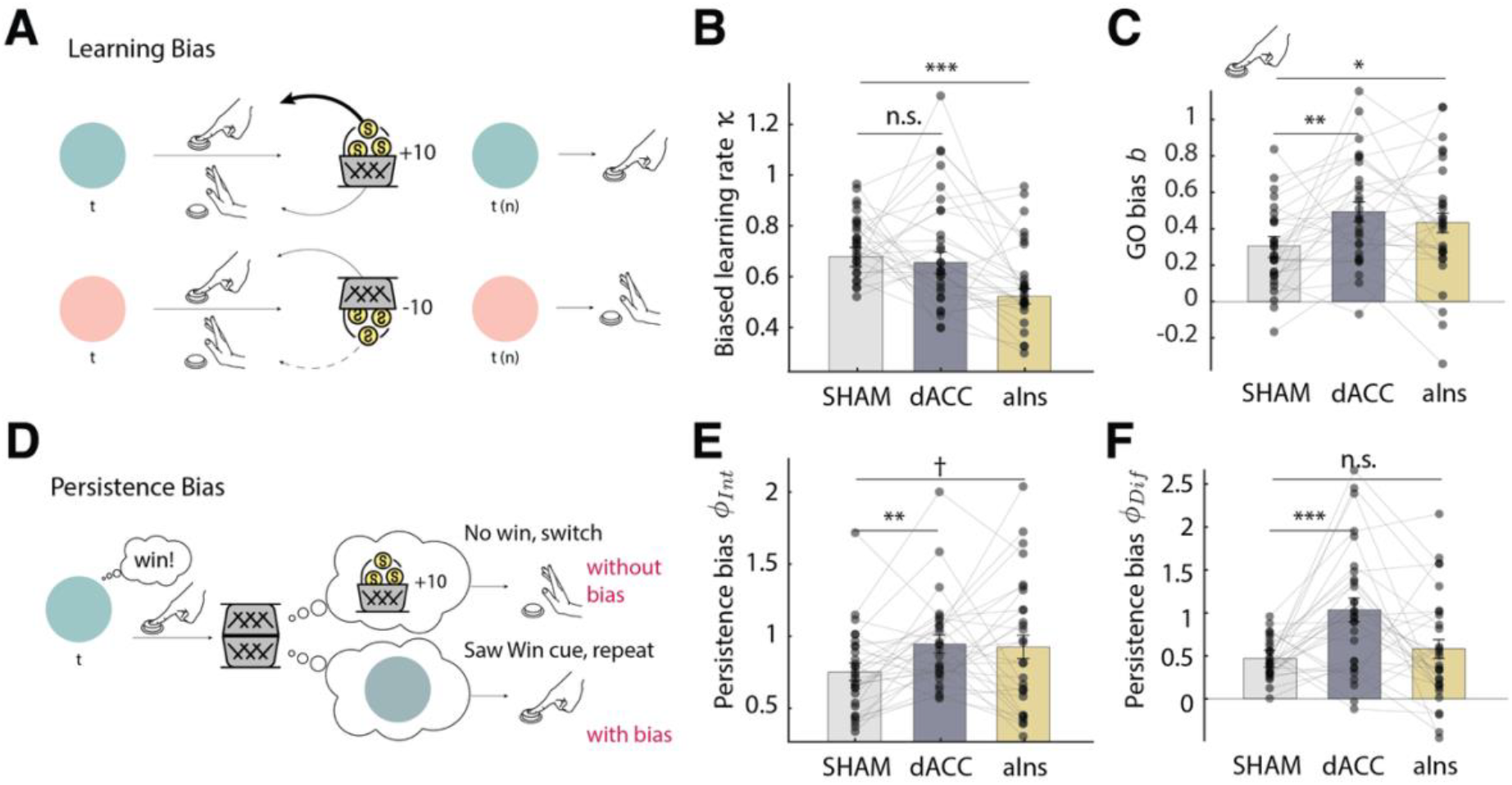
Effects of TUS on instrumental learning biases, as reflected by computational model parameters. **A**. Learning bias illustration: Learning is stronger for rewarded Go actions, and weaker for punished NoGo actions. **B**. Learning bias (κ): TUS-aIns (but not TUS-dACC) significantly diminished the learning bias, i.e., the tendency to attribute rewards to actions and the reluctance to attribute punishments to inactions. **C**. Go bias (b): both sonication conditions significantly increased participants’ overall tendency for Go responses regardless of the cue valence. **D**. Persistence bias illustration: participants show an overall higher tendency to repeat actions for Win cues than for Avoid cues. For Win cues, this implies that they sometimes ignore neutral outcomes (negative feedback, which suggests to switch the response) and instead take the Win cue itself as positive feedback (which suggests to repeat the response). **E**. Persistence parameter (φ_INT_): both sonication conditions increased participants’ overall tendency to repeat previous responses. **F**. Persistence bias (φ_DIFF_): TUS-dACC (but not TUS-aIns) significantly increased the persistence bias, leading to a stronger tendency to repeat responses relatively more for Win cues than for Avoid cues. Asterisks indicate significance levels: ^***^p < 0.001; ^**^p< 0.01; ^*^p < 0.05; † p < 0.10.

However, there was a significant difference in the learning bias κ, *F*(2, 56) = 7.680, *p* =.001, η^2^ = 0.134: this bias was significantly attenuated after TUS-aIns compared to sham, *t*(28) = 4.381, *p* <.001, Cohen’s *d* = -1.072, 95%-CI [-1.687, -0.457], but not after TUS-dACC 466 compared to sham, *t*(28) = 0.489, *p* =.629, Cohen’s *d* = -0.135, 95%-CI [-0.688, 0.419] (Fig. 4B). This observation is in line with the regression results showing an attenuated difference in repetitions of rewarded Go vs. punished NoGo responses after TUS-aIns.

Furthermore, there was a significant difference in the persistence bias φ_DIFF_, *F*(2, 56) = 9.836, *p* <.001, η^2^ = 0.157: this bias was significantly stronger after TUS-dACC compared to sham, 471 *t*(28) = 4.028, *p* <.001, Cohen’s *d* = 1.031, 95%-CI [0.397, 1.667], but not after TUS-aIns 472 compared to sham, *t*(28) = 1.058, *p* =.299, Cohen’s *d* = 0.214, 95%-CI [-0.624, 0.196] (Fig. 4F). This effect was in line with the increased cue valence effect on response repetitions vs. switches after TUS-dACC. In sum, TUS-aIns attenuated the learning bias, while TUS-dACC increased the persistence bias.

Beyond the bias parameters of interest, both TUS conditions also unspecifically increased the Go bias parameter, *F*(2, 56) = 4.958, *p* =.010, η^2^ = 0.071 (Fig. 4C), and the persistence parameter φ_INT_, *F*(2, 56) = 3.517, *p* =.036, η^2^ = 0.055 (Fig. 4E; neither significant when correcting for multiple comparisons): Relative to sham, the overall Go bias *b* was significantly increased after both TUS-dACC, *t*(28) = 3.148, *p* =.004, Cohen’s *d* = 0.720, 95%-CI [0.206, 1.233], and TUS-aIns, *t*(28) = 2.130, *p* =.042, Cohen’s *d* = 0.438, 95%-CI [0.007, 0.869]. Similarly, the overall persistence parameter φ_INT_ was significantly increased after TUS-dACC, 483 *t*(28) = 3.070, *p* =.005, Cohen’s *d* = 0.639, 95%-CI [0.181, 1.096], and marginally significantly 484 increased after TUS-aIns, *t*(28) = 1.972, *p* =.059, Cohen’s *d* = 0.429, 95%-CI [0.026, 0.884] (not significant when correcting for multiple comparisons). There was no significant difference in the feedback sensitivity parameter ρ, *F*(2, 56) = 0.928, *p* =.401, η^2^ = 0.015, or the learning rate ε, *F*(2, 56) = 1.047, *p* =.358, η^2^ = 0.018.

Taken together, the effects of dACC and aIns sonication were best captured by a decrease in the learning bias after TUS-aIns, an increase in the persistence bias after TUS-dACC, and increases in overall Go biases and persistence after both types of sonications. These effects were well captured by posterior predictive checks: Simulated learning curves closely followed the empirical learning curves (Fig. 3E, F). These observations also match the regression results of response repetitions/switches, with an attenuated difference between rewarded Go and punished NoGo responses after TUS-aIns (Fig. 3G) and a stronger cue valence effect after TUS-dACC (Fig. 3H). In sum, in this study, we found a double dissociation between the effects of TUS-aIns and TUS-dACC, with both altering different Pavlovian biases.

## Discussion

This study investigated the effects of TUS to the dACC and aIns on Pavlovian biases using a Motivational Go/NoGo task. Participants showed three forms of Pavlovian biases: a response bias, learning bias, and persistence bias (Swart et al., 2017; Algermissen et al., 2024). TUS revealed a double dissociation, with dACC and aIns affecting distinct learning processes: TUS-aIns attenuated a learning bias, reducing the tendency to attribute rewards to actions and reluctance in attributing punishments to inactions, while TUS-dACC increased a persistence bias, strengthening the tendency to use cue valence as a reinforcer signal and persist after positive (compared to negative) cues. Given that dACC and aIns are part of the same resting-state network, the saliency network (Seeley, 2019; Taylor et al., 2008), and regularly co-activate in cognitive tasks (Kurth et al., 2010; Medford and Critchley, 2010), previous research has assigned largely similar roles to these regions. Our study contributes to understanding their different roles in learning from feedback: While the dACC appear important in selecting which environmental signals to use as feedback, the aIns seems important in credit assignment whether these signals were caused by an agent’s own actions.

We set out this study under the hypothesis that perturbing neural activity in dACC and aIns, which are part of an early “reset” network that rapidly responds to negative/salient stimuli (Fouragnan et al., 2015), could alter the strength of Pavlovian biases. In particular, several previous studies have observed a selective role of dACC and aIns in choices to avoid losses (Cecchi et al., 2024; Palminteri et al., 2012), learning from negative feedback (Billeke et al., 2020; Horing and Büchel, 2022), and adjusting behaviour (McGuire and Kable, 2015; van Geen et al., 2025). However, in our results, TUS did not selectively affect responses to negative (Avoid) cues. Specifically, TUS did not change the cue valence effect (i.e., response bias), the required action effect (i.e., overall accuracy), or their interaction (i.e., a change in accuracy for only Avoid cues). Similarly, in our computational models, (baseline) learning rates and the response bias were unaffected by TUS. Instead, the effects of TUS only became apparent over time, indicative of changes in learning processes, and were spread across all four cue conditions. These findings reflect more intricate changes in learning that were well captured by a computational model (Algermissen et al., 2024). This finding might not be surprising given a broad literature implicating dACC and aIns in learning from feedback and adjusting action policies (Alexander and Brown, 2015; Behrens et al., 2007; Fouragnan et al., 2019).

We found that stimulating the aIns with TUS led to a reduction in a learning bias: participants showed a reduced tendency (i.e., lower learning rate) to ascribe rewards to their own actions, as well as a reduced reluctance to ignore punishments of their own inactions. Such biases have been interpreted as global “priors” on which action-outcome relationships likely hold in an organism’s environment (Algermissen et al., 2024; Swart et al., 2017). Using such priors, individuals can trade-off noisy outcomes of single actions against prior hypotheses on what is generally the best response strategy. Such biases will facilitate the acquisition of actions that are conducive to rewards but slow down the acquisition of actions in contexts in which punishments are involved. While such biases will usually be adaptive and lead to more robust learning, the tendency to attribute rewards to own actions can lead to the false inference of “illusionary control” over outcomes that are in fact random (Aarts et al., 2005; Ebert and Wegner, 2011). Similarly, these biases could explain why humans judge actions that lead to negative outcomes more harshly than omissions leading to such outcomes (“omission bias”) (Spranca et al., 1991), a phenomenon cited to explain some humans’ reluctance to vaccinate (Ritov and Baron, 1995, 1990).

We have previously found that neural signals associated with such biased learning are first visible in the blood-oxygen-level-dependent (BOLD) signal from cortical regions including the perigenual anterior cingulate cortex (pgACC) and posterior cingulate cortex (PCC), prior to subcortical regions (striatum) (Algermissen et al., 2024). Previous research has indeed suggested cortical regions, most prominently the dACC, to control the speed of learning (learning rates) in subcortical regions (Alexander and Brown, 2015; Behrens et al., 2007; Meder et al., 2017). Although related bias phenomena, such as auto-shaping and negative self-maintenance, have been shown in animals such as rodents or pigeons (Brown and Jenkins, 1968; Williams and Williams, 1969), we have previously speculated that a prefrontal, cortical basis of this bias in humans might hint at a more recent innovation in primates that is responsible for this particular learning bias.

Given the previous literature on the role of the dACC in scaling learning rates (Behrens et al., 2007; O’Reilly et al., 2013; Piray et al., 2019b), we would have a priori expected that TUS dACC should have affected the learning bias, and a selective reduction under TUS-aIns might at first appear puzzling. The aIns has previously been found to encode the uncertainty/risk under which a choice is made (Hsu et al., 2005; Preuschoff et al., 2008; Vilares et al., 2012) as well as the average reward rate in an environment (Wittmann et al., 2020), which are key environmental variables that should scale learning rates according to normative models (Cazé and van der Meer, 2013; Piray and Daw, 2021; Yu and Dayan, 2005). Overall, these findings suggest that aIns plays a crucial role in using higher-order beliefs about the environment to decide how much to update lower order beliefs based on reward/punishment feedback. This supports a role of the aIns in scaling learning rates, which can give rise to learning biases.

An alternative possibility is that the TUS-aIns effect reducing biased credit assignment arises from its role in social cognition and self-other attribution. Increased aIns activity has been shown both when an agent is directly affected by an outcome (e.g., pain, unpleasant taste) and when they observe such outcomes in another individual (Jabbi et al., 2008; Lamm and Singer, 2010; Singer et al., 2004). Hence, the computational challenge arises for the aIns to tell apart which outcomes were caused by oneself and which were merely observed in others (Jabbi et al., 2008). TUS-aIns might blur this boundary, leading to changes in credit assignment as observed in the current study. Some studies have speculated that this role is subserved by von Economo and Fork neurons, which, within the primate brain, occur almost exclusively in the aIns (Allman et al., 2010, 2005) and have only been observed in animal species that live in large social groups (Evrard, 2019). This speculative interpretation suggests a dedicated role of the aIns in credit assignment, following an evolutionary route that might be special in primates. Investigating the causal contribution of such deep cortical circuits to learning from feedback might greatly benefit from the use of non-invasive brain stimulation tools such as TUS. Taken together, our results suggest that aIns plays an important role in credit assignment and deciding whether environmental feedback signals were caused by personal actions.

In contrast to TUS-aIns, TUS-dACC selectively affected the persistence bias, with an increased tendency to repeat actions for Win cues than Avoid cues, regardless of the actions and outcomes they had been followed by. This bias reflects participants’ tendency to use the cue valence instead of the outcome valence as a feedback signal (Algermissen et al., 2024). Knowledge of the cue valence is important for disambiguating neutral outcomes, which can only be used as a teaching signal when contrasted against the counterfactual that could otherwise have been obtained (i.e., a neutral outcome signalling a missed reward is negative feedback; a neutral outcome signalling a missed punishment is positive feedback). However, human participants show limited ability to tell apart counterfactual and real outcomes, sometimes taking a neutral outcome obtained instead of a reward as a positive outcome, and a neutral outcome obtained instead of a punishment as a negative outcome (Algermissen et al., 2024), a tendency exacerbated by TUS-dACC. Previous studies have implicated a causal role of the dACC in learning from counterfactual feedback (Fouragnan et al., 2019) and deciding when it is worth switching an action policy (Hayden et al., 2011; Karlsson et al., 2012; Kolling et al., 2012). When perturbed with TUS, the dACC might mediate a confusion between real and counterfactual feedback, leading to a decreased ability to switch behaviour after a missed reward, which gives rise to larger persistence bias. In sum, our results suggest that the dACC is important in selecting which environmental signals to use as reinforcer to adjust behaviour.

Our findings also highlight the promise of TUS as a neuromodulation tool in cognitive neuroscience. Compared to other methods, such as TMS or transcranial direct current stimulation (tDCS), TUS allows for deeper, more focal targeting of subcortical and midline/lateralised deep cortical structures, making it particularly suited to probing the causal roles of regions like the aIns and dACC. As such, this method opens new possibilities for investigating the neural mechanisms of learning and decision-making.

In summary, our study reveals a double dissociation in the effects of TUS on learning biases, with TUS-aIns selectively attenuating the learning bias and TUS-dACC increasing the persistence bias. Hence, while dACC and aIns often co-activate in cognitive roles, they appear to have distinct roles in learning from feedback: while TUS to the aIns decreased people’s tendency to overly take credit for rewards following action and to ignore punishments following inaction, TUS to dACC increased participants’ tendency to take the cue valence as a reinforcer signal. These findings advance our understanding of the differential roles of deep cortical regions in instrumental learning and highlight the potential of TUS as a tool for modulating cognitive and affective processes. Our findings challenge the traditional view that learning biases are exclusively driven by subcortical regions associated with rigid, habit-like behaviours. Instead, these results emphasize the role of frontal inputs, which contribute to counterfactual reasoning and behavioural flexibility (Behrens et al., 2007; Fouragnan et al., 2019). The involvement of these two areas in biased learning suggests that these biases may not be rigid constraints, but rather adaptive priors that guide how individuals integrate past experiences. By balancing prior beliefs about action-outcome associations with flexible learning from rewards and punishments, these biases may enhance decision-making in dynamic environments (Algermissen et al., 2024; Algermissen and den Ouden, 2023).

## Methods

### Participants

34 healthy participants (18 females, 16 males) aged 19 to 59 years (mean age = 29.73, SD = 11.16) took part in this study. One participant was excluded because they did not understand the task and three because of missing sessions. 29 participants (16 females, 13 males), aged 19 to 59 years (mean age = 30.24, SD = 11.63) were included in the analysis. Participants had 648 normal or corrected-to-normal vision and were carefully screened for any counter indications 649 to TUS and MRI, with none reporting a current diagnosis of neurological or psychiatric disorders or use of psychoactive medications. All participants provided written informed consent following a full explanation of the study procedures. Ethical approval was granted by the University of Plymouth Faculty of Health Staff Research Ethics and Integrity Committee (reference ID: 3394; date: 28/06/22). Participants were compensated an average of £85, which included a performance bonus of up to £10 based on points collected over three sessions in the Motivational Go/NoGo task. Travel expenses were reimbursed up to £5 per session. All study sessions took place at the Brain Research & Imaging Centre (BRIC) in Plymouth, UK.

### Study design

The study design is summarised in Fig. 1A. Participants began with an MRI-only session, which included a series of scans, including a T1-weighted MRI scan. This imaging data set was used to create a personalised head model for neuronavigation and acoustic and thermal simulations, enabling precise planning of the TUS target and transducer placement for the upcoming TUS sessions. In the first session, participants completed a practice phase to become familiar with the Motivational Go/NoGo learning task.

For the TUS sessions, participants received a randomised sequence of TUS conditions. Each session involved an 80-second TUS application with three different conditions: active stimulation of the bilateral anterior insula (TUS-aIns), active stimulation targeting the dorsal anterior cingulate cortex (TUS-dACC), and a sham condition. In the sham condition, the transducer was positioned over the temporal region bilaterally, similar to the TUS-aIns setup, but no ultrasound was delivered. To maintain a consistent procedure, both sonication and sham conditions were always applied first to the left hemisphere, followed by the right.

### MRI data acquisition

For TUS planning, including neuronavigation and acoustic simulations, MRI scans were acquired using a Siemens MAGNETOM Prisma 3T scanner (syngo MR V11E, Siemens Healthineers, Erlangen, Germany) equipped with a 64-channel head coil. The scanning protocol for this study included a T1-weighted MPRAGE sequence in the sagittal plane (TR = 2100 ms, TE = 2.26 ms, inversion time = 900 ms, flip angle = 8°, GRAPPA acceleration factor 680 = 2, matrix size = 256 × 256, 176 slices, voxel size = 1 × 1 × 1 mm^3^).

### TUS protocol and procedure

Targeting of the bilateral aIns and dACC was guided by personalised anatomical landmarks using each participant’s T1-weighted MRI scans during the planning of target and transducer placement (see Supplementary Table 1). Acoustic field simulations, generated from pseudo-CT skull models derived from participants’ T1-weighted images using a convolutional neural network (CNN) (Yaakub et al., 2023), ensured precise targeting and safety by keeping both on-target and off-target pressure within acceptable ranges.

A bespoke CTX-500 NeuroFUS TPO system (Brainbox Ltd., Cardiff, UK) with a four-element annular transducer (diameter = 64 mm, central frequency = 500 kHz, and steering range between 27.3 and 82.6mm) (Supplementary Fig. 1) was used to deliver the TUS protocol. The TUS protocol consisted of repetitive TUS applied using a 5 Hz-patterned protocol with a 10% duty cycle, lasting 80 seconds and delivering a total of 400 pulses (pulse duration of 20 ms and pulse interval of 200 ms). The initial spatial-peak pulse-average intensity (ISPPA) in free-field conditions was set at 50 W/cm^2^ for all participants. We took care to remain within the guidelines for human ultrasound exposure as defined by ITRUSST (Aubry et al., 2024). The results of our acoustic simulations predominantly indicated a maximum skull temperature rise below 2°C (Supplementary Table 1). In cases where this threshold was exceeded, we calculated the CEM43, a metric reflecting both duration and intensity of heating relative to 43°C, the critical threshold for thermal cell damage. We ensured that CEM43 values remained well below 0 (Aubry et al., 2024). In our study, CEM43 was always below 0.1.

To improve ultrasound transmission, a layer of ultrasound gel (Aquasonic 100, Parker Laboratories Inc.) was applied at the transducer placement site, with a 2 cm gel pad (Aquaflex, Parker Laboratories Inc.) positioned between the transducer and the participant’s head. Participants’ heads remained unshaved, and any air bubbles were carefully eliminated by smoothing and combing the hair. Neuronavigation was performed using Brainsight v2.5 (Rogue Research Inc., Montréal, Québec, Canada) with T1-weighted anatomical MR images. During each session, focal depth measurements obtained from Brainsight were entered into the NeuroFUS TPO system (Brainbox Ltd., Cardiff, UK) before stimulation, and the trajectory 710 was sampled to support confirmatory acoustic simulations following each session.

### Blinding procedure

Sham TUS was administered identically to active TUS, except that no actual stimulation was applied. Instead, participants listened to a sound mimicking the transducer’s pulse repetition frequency through bone conduction headphones, positioned approximately 2 cm posterior and superior to the temples. This sound was only played during the Sham condition to ensure blinding, while headphones were worn in both active and Sham sessions.

### Task

The task consisted of four blocks, presented approximately 15, 25, 35, and 45 minutes after TUS (Fig. 1B, C). Each block featured four cues (gems), with each trial requiring participants to determine, through trial and error, the correct action (Go or NoGo) and the potential outcome (reward or punishment) associated with each stimulus. The task design involved four different cues that varied by cue valence (Win vs. Avoid) and required action (Go vs. NoGo), counterbalanced across participants in three sessions of 320 trials each. Motivationally incongruent cues were included, where participants’ natural action tendencies (e.g., to act for a reward) conflicted with the task requirements (e.g., to refrain from action when a reward was expected). For each cue, correct responses resulted in positive outcomes (80% chance of reward or avoiding punishment), whereas incorrect actions had only a 20% chance of yielding positive outcomes. The valence (Win or Avoid) of each cue was not explicitly signalled and had to be learned over time.

This probabilistic task was designed to capture learning and decision-making under uncertainty, similar to real-world scenarios where outcomes are not guaranteed. The structure of probabilistic feedback allowed us to observe participants’ flexibility in adjusting strategies, sensitivity to feedback, and the cognitive and neural processes underlying decisions when faced with motivational conflicts, such as balancing exploration versus exploitation in uncertain environments.

### Stimuli display

The probabilistic motivational learning task was displayed on an LCD screen (1920 × 1080 resolution, 60 Hz refresh rate, 8-bit colour depth, RGB, standard dynamic range (SDR)), positioned 0.5 m in front of participants. The experiment was programmed using the Presentation software (Neurobehavioral Systems Inc., Berkeley, CA, USA) and run on a Windows-based computer. Participants responded using their preferred hand by pressing the SPACE bar on a keyboard.

### Post-TUS questionnaire

After each TUS session, as well as on the following day, participants were asked to report any adverse effects. They were presented with a list of symptoms and asked to rate the intensity to which they experienced each of them using a 4-point scale (absent, mild, moderate, severe) and whether they thought their experience was related to the stimulation on a 5-point scale (unrelated, unlikely, possible, probable, definite). Symptoms included those reported in (Legon et al., 2020), to which we added speech problems, vision problems, and muscle tightness of the face or arm. Respondents had the possibility of listing and rating up to two additional symptoms and to provide details for their experience.

For further insights, Supplementary Table 1 present detailed information regarding the acoustic simulation parameters and the outcomes observed across participants. This includes data on mechanical indices, derated ISPPA values at the focal point, and the incidence of side effects associated with both aIns and dACC stimulation.

### Regression analyses

We fit mixed-effects logistic regression models using the lme4 package in R. We always included a random intercept per subject and all possible random slopes and random correlations, achieving a maximal random effects structure (Barr et al., 2013). We used sum-to-zero coding for all categorical predictors. We obtained *p*-values from Wald chi-square tests using Type-3 sums-of-squares via the Anova() function from the car package.

### Computational modelling and model comparison

We fit a series of increasingly complex computational reinforcement learning models to participants’ choices, following previous studies (Algermissen et al., 2024; Guitart-Masip et al., 2012; Swart et al., 2017). All models are Q-learners that learn the expected value of performing an action for a given cue using reward prediction errors and select actions with higher expected value. On a given trial t, the base model **M1** computes choice probabilities for Go and NoGo actions (a) given a cue (s) using action weights (w; modified Q-values) turned into probabilities via a softmax transform:

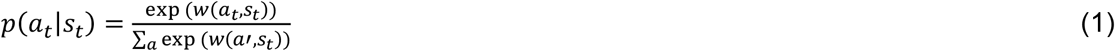

For each action, the model obtains an outcome r and computes a reward prediction error as the difference between the outcome r and the expected value Q(a, s). It then uses prediction errors to update Q-values via the delta learning rule:

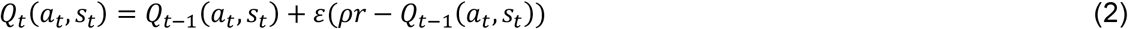

Outcomes can be +1 (reward) or 0 (no reward) for Win cues, or 0 (no punishment) or -1 (punishment) for Avoid cues. In this learning rule, prediction errors are scaled by the learning rate ε, with large learning rates leading to faster updating (and a stronger recency bias in value estimates) and small learning rates leading to slower updating (and a more long-lasting impact of outcomes obtained far in the past). Furthermore, outcomes themselves are scaled with a participant-specific feedback sensitivity parameter ρ, which plays a similar role as an inverse temperature parameter: high values of ρ lead to Q-values for Go and NoGo actions developing further apart with learning and leading to more deterministic choices, while small values of ρ lead to Q-values closer together and thus more stochastic choices. We initialized Q-values to the midpoint between the two possible outcomes of each cue (+0.5 for Win cues, -0.5 for Avoid cues) multiplied with participants’ respective feedback sensitivity parameter such that starting Q-values are the midpoint of each participant’s subjective value space. Given that the valence of each cue (Win/Avoid) was not instructed but had to be inferred from the attainment of valenced outcomes (rewards/punishments), participants could not interpret neutral outcomes as positive/negative feedback before having observed such a valence outcome. During this phase, Q-values were updated based on feedback, but “muted” in the action selection process (multiplied with zero when computing choice probabilities), assuming that participants could retrospectively infer the meaning of neutral outcomes once they determined the valence of a given cue. Upon the first encounter of a valenced outcome (reward, punishment), Q-values for this cue were “unmuted” and subsequently used for action selection.

We extended this base model using several bias terms. In **M2**, we added a Go bias term *b*to the Q-values of making a Go response for each cue, capturing participants’ overall propensity of making a Go response:

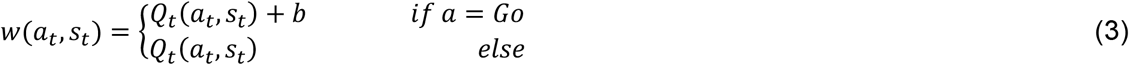

In **M3**, in addition to the Go bias, we added the response bias π multiplied with the cue valence V (+0.5 for Win; -0.5 for Avoid; value of 0.5 arbitrarily chosen for scaling) to the Q-values of making a Go response:

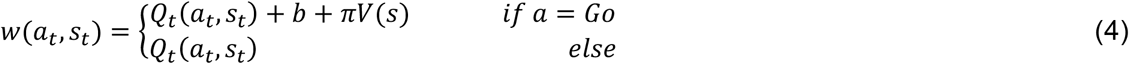

Note that the cue valence was fixed and participants instructed that each cue was either a Win or an Avoid cue. Hence, no incremental learning of the cue valence was required, but the valence could be inferred as soon the first valenced outcome was encountered. Similar to early learning from neutral outcomes, the response bias was muted (multiplied with zero) up until the first encounter of a valenced outcome and only subsequently unmuted.

In **M4**, instead of a response bias, we added a learning bias κ, capturing participants’ tendency to use an increased learning rate for rewarded Go actions (i.e., attribute rewards to their own actions) and a decreased learning rate for punished NoGo actions (i.e., an unwillingness to attribute punishments to own inactions):

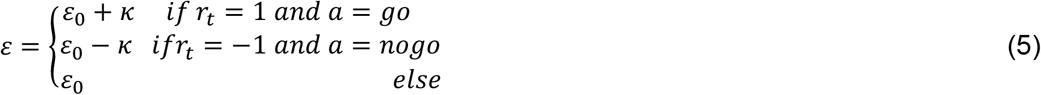

Since learning rates were sampled in a continuous space and later inverse-logit transformed to constrain them to the range of [0, 1], the impact of κ might be asymmetric after the transformation. To achieve a symmetric impact of κ, we first determined whether the base learning rate ε_0_ was smaller or bigger than 0.5, then computed one half of the bias (i.e., the side closer to the range limits), took the difference between the base and the biased learning rate, and used this difference to compute the other half of the bias:

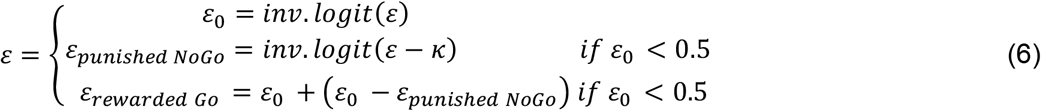

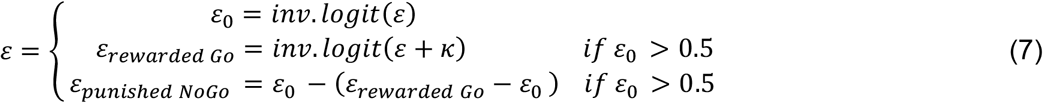

In **M5**, we added both the response bias π and the learning bias κ, which provided a better fit than each bias in isolation.

In **M6**, to capture participants’ overall tendency to repeat responses irrespective of the received outcome, we extended M5 by adding an intercept persistence parameter φ_INT_ to the weight of the action performed on the last encounter of the same cue:

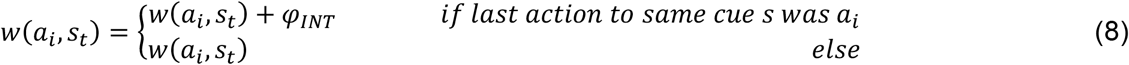

Finally, in **M7**, to capture participants’ higher tendency to repeat responses for Win than for Avoid cues, we added the persistence bias φ_DIFF_, which was added to the weight of the last performed action in case of a Win cue and subtracted from this weight in case of an Avoid cue:

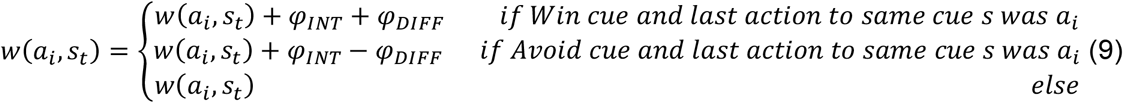

We fitted models using weekly informative hyperpriors to *X*_*ρ*_ *∽𝒩*(2, 3), *X*_*ε*_*∽𝒩*(0, 2), *X*_*b,π,κ*_ ∽*𝒩*(0, 3) in line with previous studies (Algermissen et al., 2024; Swart et al., 2017). We constrained feedback sensitivities ρ to be positive using the exponential transform and learning rates ε to the range [0, 1] using an inverse the inverse-logit transform. Furthermore, we constrained κ and φ_DIFF_ to be positive in line with the hypothesized direction of the learning and persistence biases (Fig. 2B, C) using the *y* = *log*(1 + *exp*(*x*)) transform, which is y = 0 for negative numbers, smoothly asymptotes 0 for small positive numbers, and is roughly y = x for large positive numbers. Leaving these parameters unconstrained led to the same qualitative conclusions, but a slightly inferior fit. In line with recent recommendations (Valton et al., 2020), we fitted the data of each sonication condition separately.

We fitted all models using the CBM toolbox in MATLAB (Piray et al., 2019a) using three steps: In the first step, we fitted the respective model to each participants’ data individually using an iterative expectation-maximization algorithm that uses the Laplacian approximation for estimating the parameter posteriors. Chances of getting stuck in a local minimum were mitigated by using multiple parameter initialisations. In a second step, we used the individual fits of all participants as the starting point in a hierarchical Bayesian inference procedure. Group-level parameters were iteratively fitted using mean-field variational Bayes and served as hyperpriors for the individual participants’ parameters. Thus, the parameter values of each participant were constrained/informed by the parameter values of all other participants, leading to more robust estimates and a lower chance of overfitting. We used these parameter values for all our posterior predictive checks. Finally, in a third step, we fitted all candidate models for all participants in one single step by iteratively (a) determining the probability that a given participant X was best fitted by a given model Y and (b) fitting the group-level parameters based on (only) those participants whose behaviour was in fact best characterized by model Y (i.e., parameters weighted proportionally to participants’ latent “model responsibility” weights). This approach allows for simultaneous hierarchical Bayesian inference (with participants constraining each other’s parameter values) and random-effects model selection (allowing for the possibility that different participants are best characterized by different models). Based on this last step, we computed the model frequency and protected exceedance probability of each model and performed Bayesian model selection (Stephan et al., 2009), selecting the model with the highest exceedance probability as the winning model.

We compared parameters across conditions using one-way repeated-measures ANOVAs as implemented in the ezAnova() function of the ez package in R. We computed paired *t*-tests using the t.test() function from the stats package in R and computed paired-samples Cohen’s *d* using the Cohen.d() function from the effsize package in R.

To validate that our best fitting model M7 could reproduce key patterns observed in the empirical data, we performed posterior predictive checks (Nassar and Frank, 2016; Palminteri et al., 2017) using one-step-ahead predictions (Steingroever et al., 2014). We used each participant’s best fitting parameters from the hierarchical Bayesian fit and their actual choices and outcomes from the empirical data to simulate the action probabilities of each participant on each trial, which we used as synthetic data to then plot the proportions of Go response/response repetitions the same way as the empirical data.

To confirm that we could reliably measure individual differences in model parameters, we performed parameter recovery (Algermissen and den Ouden, 2024b; Wilson and Collins, 2019). We first fitted a multivariate normal distribution to the best fitting parameter values of M7, sampled 1,000 new combinations of parameter values from this distribution, simulated new data based on this parameter combinations, and fitted M7 to each simulated data set. We then correlated the sampled “ground-truth” parameters to the fitted “recovered” parameters. To test whether these correlations were significantly higher than expectable by chance, we compared them against a permutation null distribution obtained by randomly permuting the assignment of ground-truth to recovered parameter values 10,000 times and computing the 95^th^ percentile of this distribution as the upper bound of a one-sided confidence interval.

To confirm that we could reliably detect the best fitting model for a given data set, we performed model recovery (Algermissen and den Ouden, 2024b; Wilson and Collins, 2019). For each of the seven candidate models, we fitted a multivariate normal distribution to the best fitting parameter values and then sampled 1,000 new parameter value combinations for each model (in total 7,000 parameter sets). We applied the following parameter constraints: ρ < 400 (otherwise possibility of infinitely large numbers), ε > 0.05 (otherwise too little learning), and ρ, π, κ, φ_INT_ and φ_DIFF_ being far enough away from zero (discarding the lowest 10% of absolute sampled parameter values). These constraints were important to ensure that each data set expressed characteristics of the respective generating model; otherwise, if one of the bias parameters was too close to zero, a more complex model would effectively reduce to a simpler model. We then simulated a new data set for each parameter combination (in total 7,000 simulated data sets) and fitted each of the seven models to each simulated data set (in total 49,000 fits). For each data set, we used the log model evidence to determine which model fitted the data best. We then computed the forward confusion matrix with the conditional probabilities (relative frequencies) of model Y emerging as a best fitting model for a data set generated by model X. We also computed the inverse confusion matrix with the conditional probabilities of model X being the generative model for a data set best fitted by model Y. To test whether the on-diagonal probabilities of these matrices were significantly higher than expectable by chance, we created a permutation null distribution by randomly permuting the log model evidences for the different fits to a given data set, computing the respective confusion matrices, and saving the on-diagonal probabilities. We again computed the 95^th^ percentile of these distributions as upper bounds of one-sided confidence intervals.

## Data availability

The data files are available as.csv and.mat files on the OSF repository https://osf.io/pux6s/?view_only=c7dab129b4f246ed937e16e5941f04da.

## Code availability statement

Analyses code to reproduce the presented acoustic simulations as well as behavioural, RS-fMRI, MRS, and self-report questionnaire results are in a GitHub repository under https://github.com/johalgermissen/mgng_tus_dacc_ains.

## Competing interests

All authors declare that they have no conflicts of interest to disclose.

## Acknowledgement

Nomi Koutsoumpari has received pump priming funds from the School of Psychology, and from the Bain Research Imaging Center, University of Plymouth for this study. Elsa Fouragnan is funded by a UKRI FLF (MR/Y034368/1), a BBSRC (BB/Y001494/1), a Neuromod+ grant (EP/W035057/1) and an ARIA grant (SCNI-PR01-P15).

## Notes

### Competing Interest Statement

The authors have declared no competing interest.

### Summary of Updates

We rephrased sections of the Introduction and Discussion for clarity and conciseness. In Figure 1A, we updated the depiction of the brain by adding an overlay of the T1 image along with averages of pressure and temperature. Minor wording adjustments were made throughout the manuscript. Additionally, the term perseveration bias was replaced with persistence bias for greater accuracy and consistency.

