## Supplementary Material for "Ultrasound neuromodulation reveals distinct roles of the dorsal anterior cingulate cortex and anterior insula in learning"

### Supplementary Results

We investigated the impact of TUS on Pavlovian biases in both response execution and learning. Using regression models, we analyzed participants' Go/NoGo choices along with their tendencies to repeat or switch responses. This analysis revealed a significant 4-way interaction between required action, cue valence, sonication condition, and block half, $\chi^2(2) = 21.548$, $p < .001$. To further understand these effects, we analysed each cue condition (Go-to-Win, Go-to-Avoid, NoGo-to-Win, NoGo-to-Avoid) separately.

For Go-to-Win cues, there was a significant main effect of block half, $b = 1.051$, 95%-CI [0.788, 1.314], $\chi^2(1) = 61.484$, $p < .001$, with the proportion of Go responses increasing over time, reflecting learning. The main effect of sonication condition was not significant, $\chi^2(2) = 1.727$, $p = .422$, with no overall difference in responding between TUS-dACC and sham, $b = -0.182$, 95%CI [-0.587, 0.222], $\chi^2(1) = 0.779$, $p = .377$, or between TUS-aIns and sham, $b = -0.244$, 95%-CI [0.671, 0.183], $\chi^2(1) = 1.252$, $p = .263$. In contrast, the interaction between block half and sonication was significant, $\chi^2(2) = 16.003$, $p < .001$. The increase in Go responses was attenuated after TUS relative to sham for both sonication sites (dACC vs. Sham x block half: $b = -0.231$, 95%CI [-0.425, -0.038], $\chi^2(1) = 5.485$, $p = .019$;TUS-aIns x sham x block half: $b = -0.363$, 95%-CI [0.537, -0.188], $\chi^2(1) = 16.534$, $p < .001$). Together, learning of Go responses for Win cues was attenuated after both TUS-dACC and TUS-aIns.

For Go-to-Avoid cues, there was as well a significant main effect of block half, $b$ = 1.023, 95%-CI [0.691, 1.355], $\chi^2(1)$ = 36.525, $p$ < .001, with the proportion of Go responses increasing over time, reflecting learning. The main effect of sonication was not significant, $\chi^2(2)$ = 4.337, $p$ = .114, with no significant difference in responding between sham and TUS-dACC, $b$ = -0.009, 95%-CI [-0.279, 0.261], $\chi^2(1)$ = 0.004, $p$ = .950, but a marginally significant increase in responding after TUS-aIns compared to sham, $b$ = 0.286, 95%-CI [-0.025, 0.597], $\chi^2(1)$ = 3.256, $p$ = .071. The interaction between block half and sonication was not significant, $\chi^2(2)$ = 2.418, $p$ = .299, with no difference in the effect of block half between TUS-dACC and sham, $b$ = -0.003, 95%-CI [-0.191, 0.185], $\chi^2(1)$ = 0.001, $p$ = .978, or between TUS-aIns and sham, $b$ = 0.139, 95%-CI [-0.081, 0.359], $\chi^2(1)$ = 1.538, $p$ = .215. Taken together, there was some evidence for overall more Go responses to Goto-Avoid cues after TUS-aIns, but this effect was only marginally significant. There was no significant evidence that this effect became stronger or weaker with learning after either sonication condition.

For NoGo-to-Win cues, there was a significant main effect of block half, $b$ = -0.836, 95%-CI [1.090, -0.582], $\chi^2(1)$ = 41.533, $p$ < .001, with the proportion of Go responses decreasing over time, reflecting learning. The main effect of sonication was not significant, $\chi^2(2)$ = 3.328, $p$ = .189, with only a marginally significant increase in responding after TUS-dACC compared to sham, $b$ = 0.288, 95%-CI [-0.040, 0.615], $\chi^2(1)$ = 2.955, $p$ = .086, and no significant increase in responding after TUS-aIns compared to sham, $b$ = 0.274, 95%-CI [-0.081, 0.629], $\chi^2(1)$ = 2.287, $p$ = .130. The interaction between block half and sonication was not significant, $\chi^2(2)$ = 2.298, $p$ = .317, with no difference in the effect of block half between TUS-dACC and sham, $b$ = -0.109, 95%-CI [-0.305, 0.087], $\chi^2(1)$ = 1.196, $p$ = .274, but a marginally significantly weaker effect of block half after TUSaIns compared to sham, $b$ = 0.168, 95%-CI [-0.013, 0.350], $\chi^2(1)$ = 3.296, $p$ = .069. Taken together, there was some evidence for overall more Go responses to NoGo-to-Win cues after TUS-dACC, and a weaker learning effect for such cues after TUS-aIns, but both effects were only marginally significant.

For NoGo-to-Avoid cues, there was a significant main effect of block half, $b$ = -0.963, 95%-CI [1.149, -0.195], $\chi^2(1)$ = 102.403, $p$ < .001, with the proportion of Go responses decreasing over time, reflecting learning. The main effect of sonication was not significant, $\chi^2(2)$ = 2.927, $p$ = .232, with no significant difference in responding between TUS-dACC and sham, $b$ = 0.120, 95%-CI [0.088, 0.329], $\chi^2(1)$ = 1.282, $p$ = .258, and only a marginally significant increase in responding after TUS-aIns compared to sham, $b$ = 0.152, 95%-CI [-0.027, 0.331], $\chi^2(1)$ = 2.770, $p$ = .096. The interaction between block half and sonication was not significant, $\chi^2(2)$ = 1.570, $p$ = .456, with no difference in the effect of block half between TUS-dACC and sham, $b$ = -0.023, 95%-CI [-0.182, 0.135], $\chi^2(1)$ = 0.084, $p$ = .772, or between TUS-aIns and sham, $b$ = -0.052, 95%-CI [-0.193, 0.089], $\chi^2(1)$ = 0.523, $p$ = .470. Taken together, there was some evidence for overall more Go responses to NoGo-to-Avoid cues after TUS-aIns, but this effect was only marginally significant.

In sum, there were significant effects of both TUS-dACC and TUS-aIns on Go/NoGo responses. The two active sonication conditions affected responses in all cue conditions. While some effects occurred in both the first and second half of each block, other were stronger in the second half, suggestive that the sonications affected learning. However, the effects on each cue condition taken in isolation were only significant for Go-to-Win conditions and only marginally significant for the other conditions. To better understand these effects, we followed them up with the regression analyses of response repetitions vs. switches (see main text).

**Supplementary Figure 1**

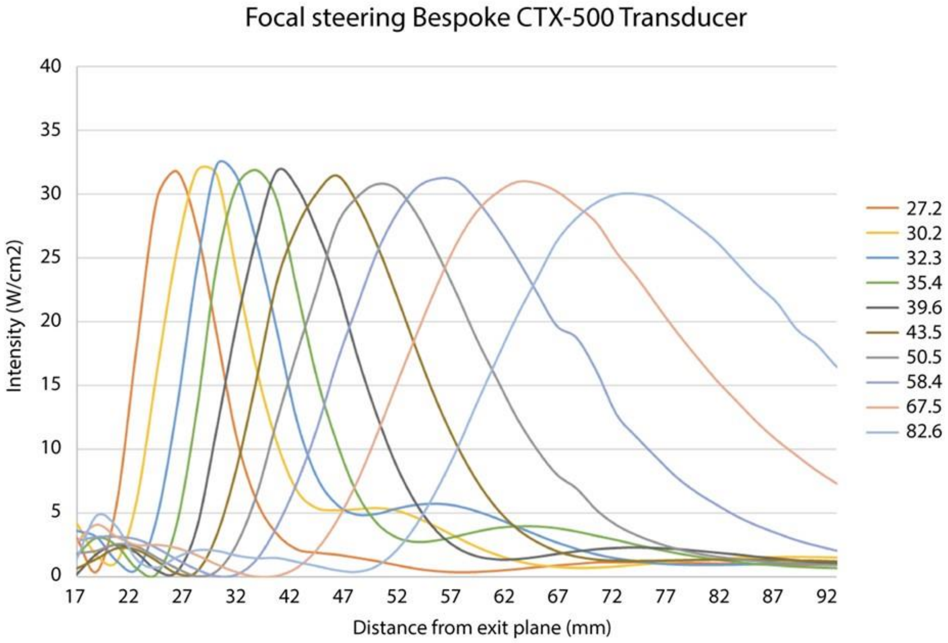

**Suppl. Fig. 1. Free-field acoustic simulations were conducted at an ISPPA of 30 W/cm².** As the focus is steered axially, the intensity naturally decreases as the distance from the coherent focus increases. The TPO compensates for this decrease by adjusting the power, ensuring a consistent intensity level. Therefore, along the steering range from 27.3 mm to 82.6 mm (measured from the transducer's exit plane), the power is regulated to maintain a constant ISPPA. The calibrated axial intensity profile plots are displayed in the figure at nine positions within the focal steering range from 27.3 mm to 82.6 mm. The figure is extracted from the manufacturer report.

**Supplementary Figure 2**

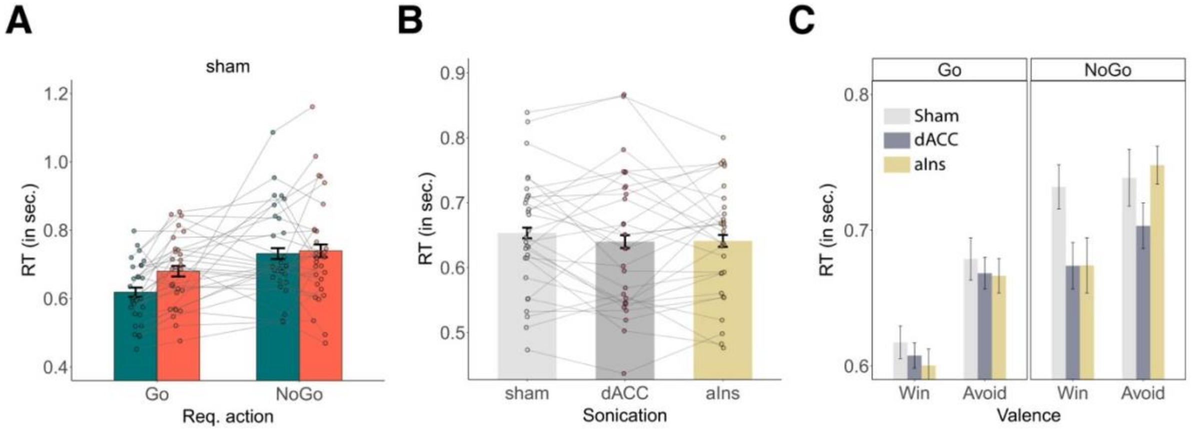

**Suppl. Fig. 2. A.** RTs for TUS-Sham split by required action (equivalent to accuracy given that RTs are only available for Go responses, and Go responses to Go cues are correct, Go responses to NoGo cues are incorrect) and cue valence. Participants showed faster RTs for (correct) responses to Go cues than for (incorrect) responses to NoGo cues, *b* = -0.207, 95% CI [-0.301 0.113], $\chi^2(1) = 18.505$, *p* < .001 and for responses to Win than to Avoid cues, *b* = -0.081, 95% CI [-0.141 -0.021], $\chi^2(1) = 6.927$, *p* = .008, reflecting a Pavlovian response bias also in RTs. This bias was slightly stronger for Go than NoGo cues, *b* = -0.061, 95% CI [-0.107 -0.014], $\chi^2(1) = 6.583$, *p* = .010. **C**. RTs per sonication condition (Sham dACC, aIns). There were no differences in RTs between conditions, $\chi^2(2) = 1.613$, *p* = .446. **C**. RTs split by required action and valence for all three sonication conditions. The effect of required action on RTs was significantly stronger after TUS-dACC compared to sham, *b* = -0.055, 95% CI [-0.099 -0.010], $\chi^2(1) = 5.773$, *p* = .016, with slower errors (responses to NoGo cues) after TUS-dACC. Furthermore, the effect of cue valence on RTs was significantly stronger after TUS-aIns compared to sham, *b* = 0.037, 95% CI [0.002 0.071], $\chi^2(1) = 4.404$, *p* = .036, driven by faster responses to Win cues after TUS-aIns.

**Supplementary Figure 3**

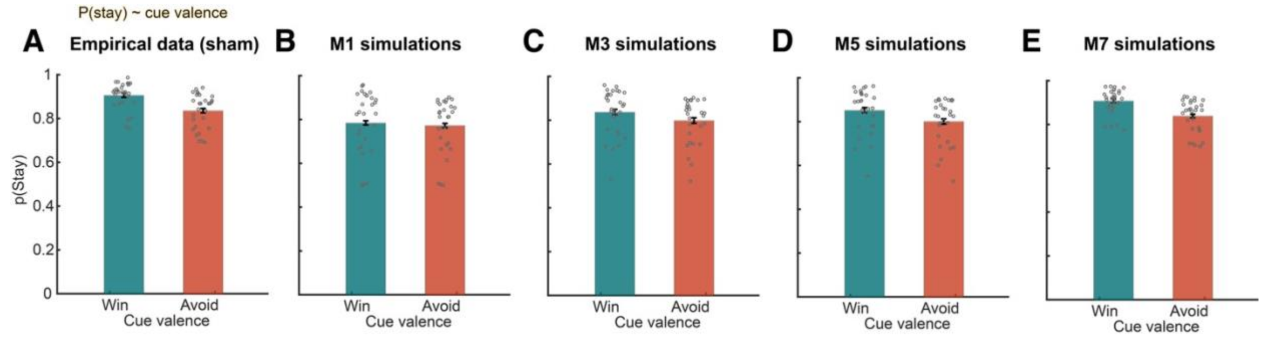

**Suppl. Fig. 3. Pavlovian learning bias illustration for sham condition. A.** Probability of response repetitions given the previous outcome (reward vs. punishment; neutral outcomes omitted in this figure) and the previously performed action (Go vs. NoGo) across empirical data and models M1, M3, M5, and M7 in the sham dataset **B-E.** The plot illustrates the Pavlovian learning bias by demonstrating how the probability of the effect of outcome valence on future response repetitions is stronger for Go than for NoGo responses, which is captured in M5 and M7 which feature a learning bias, but not in simpler models (M1, M3) without such a bias.

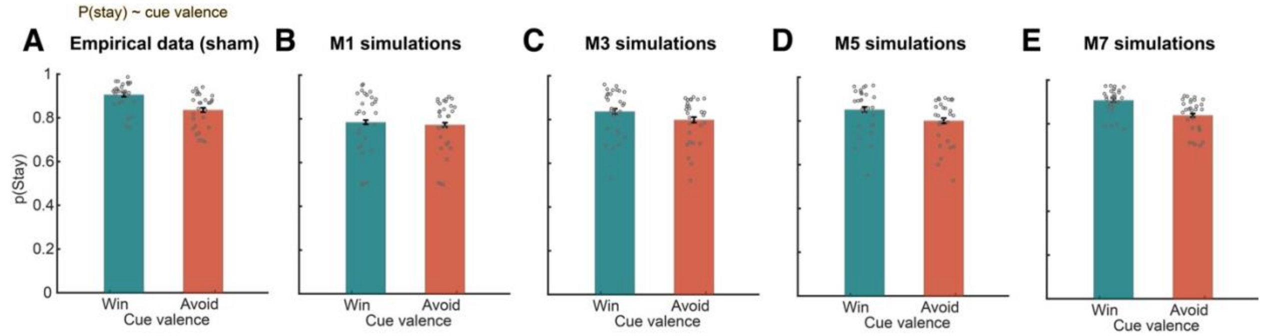

**Supplementary Figure 4**

**Suppl. Fig. 4. Persistence bias Illustration for Sham condition. A.** Probability of response repetitions given the cue valence (Win vs. Avoid) across all trials only in the sham dataset. Participants show more response repetitions for Win than Avoid cues, a tendency that is sufficiently captured by a model featuring a persistence bias (M7) seen in panel E, but less so or not at all by simpler models (M1, M3, M5), in panels B-D.

**Supplementary figure 5**

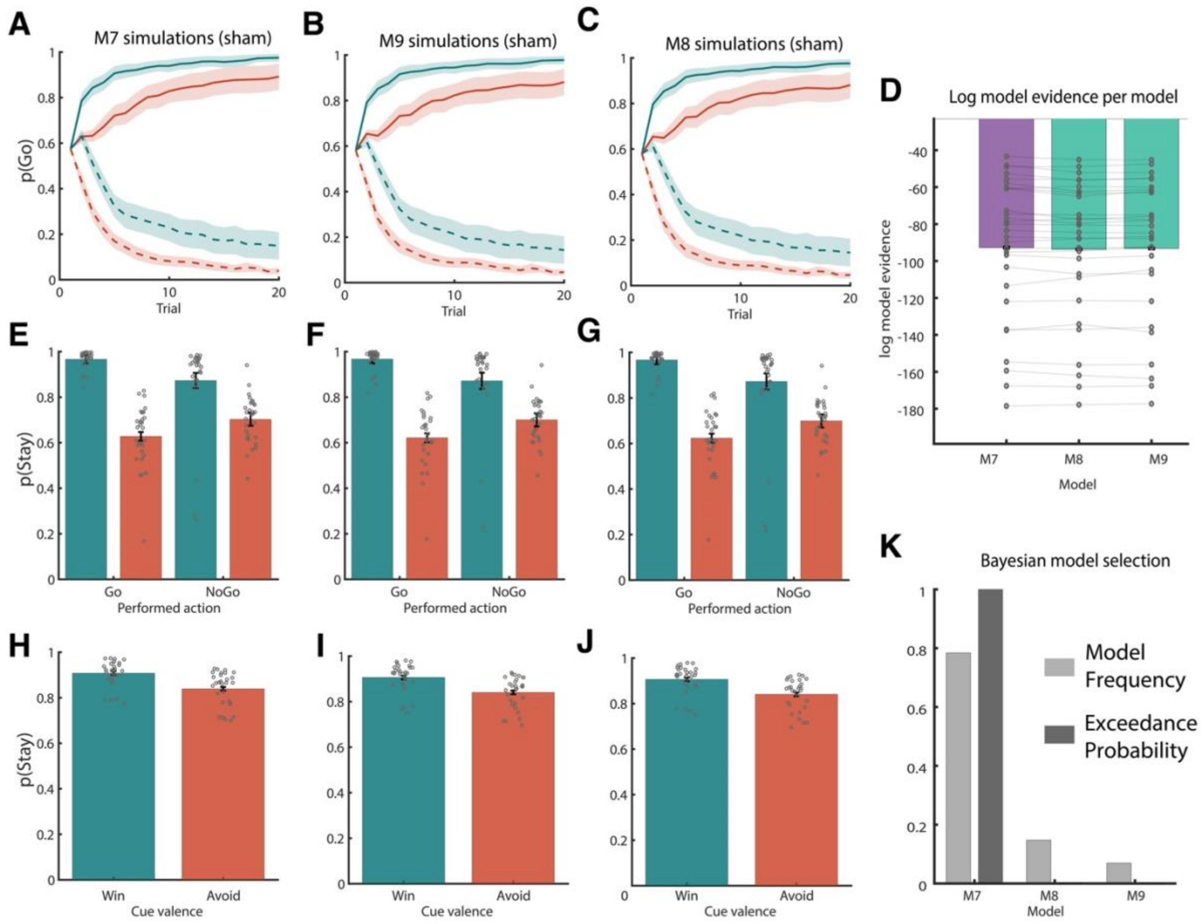

**Suppl. Fig. 5. Control models for the cue valence-specific** persistence **bias. A.** Simulations are shown for Model M7 (winning model in the main text) in which cue valence directly modulates persistence, **B.** M8, in which cue valence is added as a modulatory term to the prediction errors for trials with neutral outcomes, and **C.** M9, in which cue valence as added to the prediction errors for all trials. All models make very similar qualitative predictions regarding p(Go) per cue type, regarding p(repeat) after reward Go/punished NoGo actions (panels **D-F**), and p(repeat) for Win/Avoid cues (panels **G-I**). Still, both the log-model evidence (panel **J**). **K.** Bayesian model selection clearly favours M7 as the best model, suggesting that the behavioural finding of higher persistence for Win than Avoid cues is best captured as a modulation of the constant persistence term rather than a modulation of the prediction error term.

**Supplementary Figure 6**

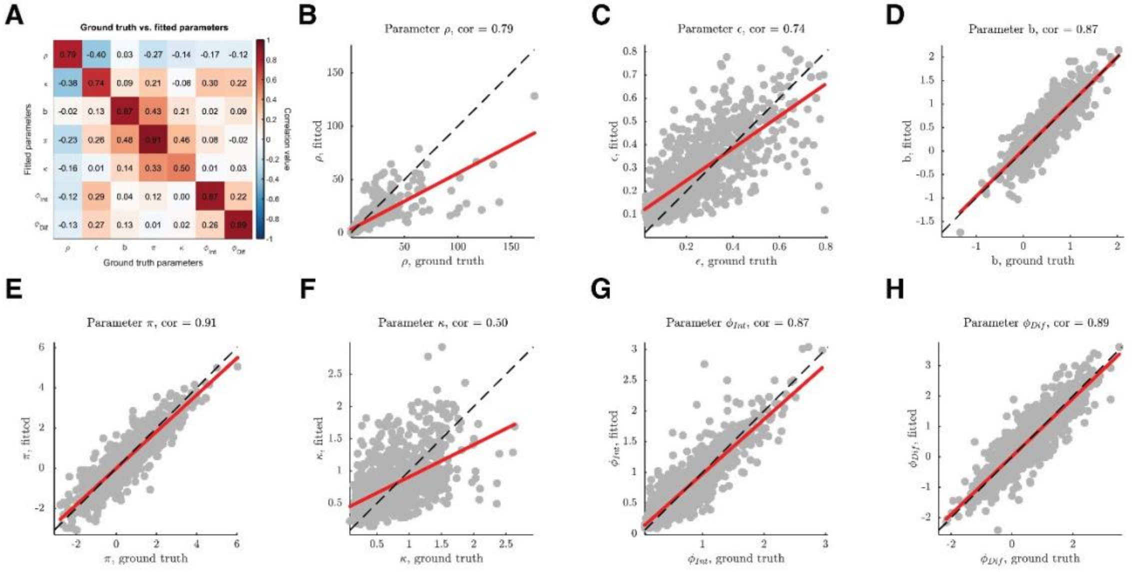

**Suppl. Fig. 6 Parameter recovery.** When simulating 1,000 new data sets from ground-truth parameters and fitting model M7 to these data sets, the fitted parameters correlate highly with the ground truth parameters, demonstrating the ability of the model to reliably capture individual differences in parameters. **A.** Heatmap of correlations between ground truth (x-axis) and fitted (yaxis) model parameters. Parameters were well recoverable (on-diagonal correlations: range 0.50–0.91, median 0.86) with only small off-diagonal correlations (all < |0.43|). All on-diagonal correlations were significantly higher than expectable under a permutation null distribution (1,000 permutations; 95$^{th}$ percentile: 0.081). **B–H.** On-diagonal correlations for the feedback sensitivity (ρ), learning rate (ϵ), Go bias (b), Pavlovian response bias (π), Pavlovian learning bias (κ), persistence parameter ($\varphi_{INT}$), persistence bias ($\varphi_{DIFF}$) from simulated data.

**Supplementary Figure 7**

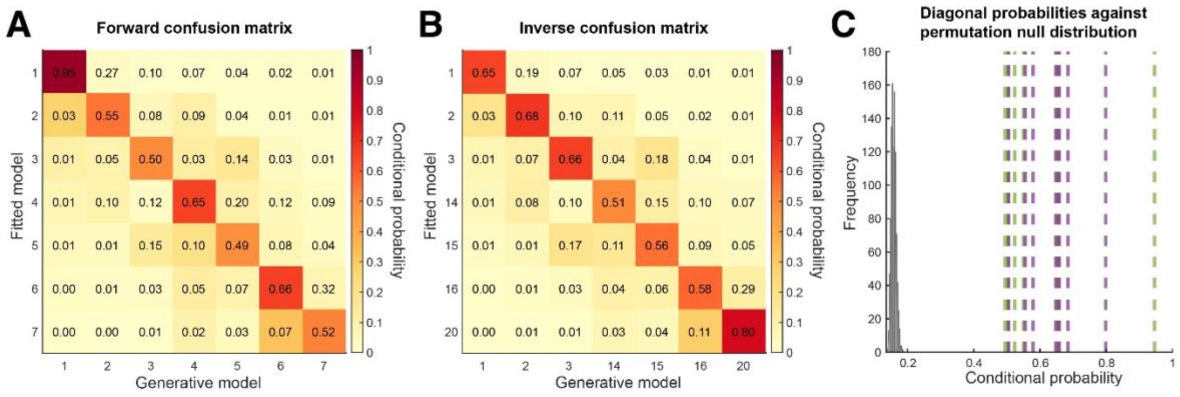

**Suppl. Fig. 7. Model recovery.** When simulating 1,000 new data sets from each model, fitting each data set with each model under consideration, and determining the best fitting model for each data set, the best fitting model most often corresponds to the original generative model, demonstrating the ability of reliably distinguish different models based on the data used in this experiment. **A.** Forward confusion matrix: A heatmap showing the conditional probability that data generated by a given model X (x-axis) is best fitted by model Y (y-axis). The diagonal elements show the probability of reidentifying the original generative model. All these probabilities are significantly higher than expected under a permutation null distribution (range 0.49–0.95, median 0.55; 95$^{th}$ percentile of permutation null distribution with 1,000 permutations: 0.171). **B.** Inverse confusion matrix: A heatmap displaying the probability that a data set best fitted by a given model X (x-axis) was in fact generated by model Y (y-axis). The diagonal elements show the probability that the best fitting model is indeed the original generative model. All these probabilities are significantly higher than expected under a permutation null distribution (range 0.64–0.80, median 0.65; 95$^{th}$ percentile of permutation null distribution with 1,000 permutations: 0.171). **C.** Diagonal probabilities vs. permutation null distribution: The histogram displays the expectable on-diagonal conditional probabilities under a permutation null distribution (grey). Dashed vertical lines display the diagonal probabilities observed in the empirical confusion matrices (purple for inverse confusion matrix, green for forward confusion matrix), which are all higher than expectable under the null distribution.

**Supplementary Table 1**

| ID | | | Temperature | Pressure | ISPPA | Transcranial Mechanical Index (MItc) |
|---|---|---|---|---|---|---|
| **sub-1** | **l-aIns** | **Skull** | 38.49 | 243.37 | 1.96 | 0.34 |
| | | **Soft Tissue** | 37.83 | 422.41 | 5.91 | 0.6 |
| | | **Sphere** | 37.15 | 422.41 | 5.91 | 0.6 |
| | **dACC** | **Skull** | 38.39 | 273.41 | 2.45 | 0.39 |
| | | **Soft Tissue** | 37.77 | 432.22 | 6.19 | 0.61 |
| | | **Sphere** | 37.16 | 432.22 | 6.19 | 0.61 |
| | **r-aIns** | **Skull** | 38.77 | 282.98 | 2.64 | 0.4 |
| | | **Soft Tissue** | 37.9 | 417.45 | 5.76 | 0.59 |

| | | | | | | |
|---|---|---|---|---|---|---|
| | | **Sphere** | 37.14 | 417.45 | 5.76 | 0.59 |
| **sub-2** | **l-aIns** | **Skull** | 38.62 | 292.45 | 2.83 | 0.41 |
| | | **Soft Tissue** | 38.03 | 618.4 | 12.56 | 0.87 |
| | | **Sphere** | 37.23 | 618.4 | 12.56 | 0.87 |
| | **dACC** | **Skull** | 38.69 | 268.83 | 2.38 | 0.38 |
| | | **Soft Tissue** | 38 | 435.65 | 6.28 | 0.62 |
| | | **Sphere** | 37.16 | 435.65 | 6.28 | 0.62 |
| | **r-aIns** | **Skull** | 39.36 | 360.88 | 4.31 | 0.51 |
| | | **Soft Tissue** | 38.61 | 527.96 | 9.2 | 0.75 |
| | | **Sphere** | 37.23 | 527.96 | 9.2 | 0.75 |
| **sub-3** | **l-aIns** | **Skull** | 38.92 | 332.94 | 3.66 | 0.47 |
| | | **Soft Tissue** | 38.16 | 530.7 | 9.31 | 0.75 |
| | | **Sphere** | 37.21 | 530.7 | 9.31 | 0.75 |
| | **dACC** | **Skull** | 38.87 | 311.74 | 3.2 | 0.44 |
| | | **Soft Tissue** | 37.97 | 471.7 | 7.36 | 0.67 |
| | | **Sphere** | 37.19 | 471.7 | 7.36 | 0.67 |
| | **r-aIns** | **Skull** | 40.04 | 444.88 | 6.48 | 0.63 |
| | | **Soft Tissue** | 38.94 | 463 | 7.09 | 0.65 |
| | | **Sphere** | 37.24 | 463 | 7.09 | 0.65 |
| **sub-4** | **l-aIns** | **Skull** | 38.64 | 300.33 | 2.99 | 0.42 |
| | | **Soft Tissue** | 37.96 | 503.14 | 8.38 | 0.71 |
| | | **Sphere** | 37.2 | 503.14 | 8.38 | 0.71 |

| | | | | | | |
|---|---|---|---|---|---|---|
| | **dACC** | **Skull** | 38.99 | 336.52 | 3.75 | 0.48 |
| | | **Soft Tissue** | 37.99 | 454.44 | 6.84 | 0.64 |
| | | **Sphere** | 37.18 | 454.44 | 6.84 | 0.64 |
| | **r-aIns** | **Skull** | 39.71 | 416.47 | 5.71 | 0.59 |
| | | **Soft Tissue** | 38.57 | 509.83 | 8.57 | 0.72 |
| | | **Sphere** | 37.21 | 509.83 | 8.57 | 0.72 |
| **sub-5** | **l-aIns** | **Skull** | 38.83 | 295.53 | 2.89 | 0.42 |
| | | **Soft Tissue** | 38.06 | 554.19 | 10.16 | 0.78 |
| | | **Sphere** | 37.24 | 554.19 | 10.16 | 0.78 |
| | **dACC** | **Skull** | 38.42 | 285.43 | 2.66 | 0.4 |
| | | **Soft Tissue** | 37.75 | 310.88 | 3.2 | 0.44 |
| | | **Sphere** | 37.09 | 310.88 | 3.2 | 0.44 |
| | **r-aIns** | **Skull** | 38.46 | 284.13 | 2.67 | 0.4 |
| | | **Soft Tissue** | 37.84 | 540.6 | 9.66 | 0.76 |
| | | **Sphere** | 37.22 | 540.6 | 9.66 | 0.76 |
| **sub-6** | **l-aIns** | **Skull** | 39.45 | 341.65 | 3.87 | 0.48 |
| | | **Soft Tissue** | 38.37 | 408.74 | 5.54 | 0.58 |
| | | **Sphere** | 37.15 | 408.74 | 5.54 | 0.58 |
| | **dACC** | **Skull** | 38.26 | 242.77 | 1.93 | 0.34 |
| | | **Soft Tissue** | 37.67 | 461.82 | 7.06 | 0.65 |
| | | **Sphere** | 37.18 | 461.82 | 7.06 | 0.65 |
| | **r-aIns** | **Skull** | 39.21 | 359.75 | 4.29 | 0.51 |

| | | | | | | |
|---|---|---|---|---|---|---|
| | | **Soft Tissue** | 38.25 | 437.08 | 6.33 | 0.62 |
| | | **Sphere** | 37.16 | 437.08 | 6.33 | 0.62 |
| **sub-7** | **l-aIns** | **Skull** | 39.26 | 316.29 | 3.32 | 0.45 |
| | | **Soft Tissue** | 38.25 | 433.7 | 6.22 | 0.61 |
| | | **Sphere** | 37.15 | 433.7 | 6.22 | 0.61 |
| | **dACC** | **Skull** | 38.85 | 330.79 | 3.58 | 0.47 |
| | | **Soft Tissue** | 38.1 | 430.84 | 6.15 | 0.61 |
| | | **Sphere** | 37.17 | 430.84 | 6.15 | 0.61 |
| | **r-aIns** | **Skull** | 38.61 | 269.4 | 2.41 | 0.38 |
| | | **Soft Tissue** | 37.96 | 490.88 | 7.97 | 0.69 |
| | | **Sphere** | 37.2 | 490.88 | 7.97 | 0.69 |
| **sub-8** | **l-aIns** | **Skull** | 38.76 | 283.16 | 2.6 | 0.4 |
| | | **Soft Tissue** | 38.04 | 477.24 | 7.53 | 0.67 |
| | | **Sphere** | 37.18 | 477.24 | 7.53 | 0.67 |
| | **dACC** | **Skull** | 38.92 | 356.12 | 4.12 | 0.5 |
| | | **Soft Tissue** | 37.99 | 447.94 | 6.65 | 0.63 |
| | | **Sphere** | 37.18 | 447.94 | 6.65 | 0.63 |
| | **r-aIns** | **Skull** | 38.84 | 288.84 | 2.75 | 0.41 |
| | | **Soft Tissue** | 38.05 | 462.33 | 7.08 | 0.65 |
| | | **Sphere** | 37.18 | 462.33 | 7.08 | 0.65 |
| **sub-9** | **l-aIns** | **Skull** | 39.01 | 318.73 | 3.36 | 0.45 |

| | | | | | | |
|---|---|---|---|---|---|---|
| | | **Soft Tissue** | 38.19 | 524.43 | 9.09 | 0.74 |
| | | **Sphere** | 37.22 | 524.43 | 9.09 | 0.74 |
| | **dACC** | **Skull** | 38.68 | 358.95 | 4.27 | 0.51 |
| | | **Soft Tissue** | 38.05 | 403.28 | 5.4 | 0.57 |
| | | **Sphere** | 37.16 | 403.28 | 5.4 | 0.57 |
| | **r-aIns** | **Skull** | 40.01 | 422.82 | 5.89 | 0.6 |
| | | **Soft Tissue** | 38.79 | 513.15 | 8.71 | 0.73 |
| | | **Sphere** | 37.22 | 513.15 | 8.71 | 0.73 |
| **sub-10** | **l-aIns** | **Skull** | 38.73 | 333.48 | 3.65 | 0.47 |
| | | **Soft Tissue** | 38.03 | 490.19 | 7.95 | 0.69 |
| | | **Sphere** | 37.2 | 490.19 | 7.95 | 0.69 |
| | **dACC** | **Skull** | 39.06 | 326.93 | 3.53 | 0.46 |
| | | **Soft Tissue** | 38.12 | 463.82 | 7.14 | 0.66 |
| | | **Sphere** | 37.19 | 463.82 | 7.14 | 0.66 |
| | **r-aIns** | **Skull** | 38.97 | 312.09 | 3.22 | 0.44 |
| | | **Soft Tissue** | 38.1 | 469.57 | 7.28 | 0.66 |
| | | **Sphere** | 37.18 | 469.57 | 7.28 | 0.66 |
| **sub-11** | **l-aIns** | **Skull** | 38.36 | 270.07 | 2.4 | 0.38 |
| | | **Soft Tissue** | 37.83 | 483.66 | 7.75 | 0.68 |
| | | **Sphere** | 37.2 | 483.66 | 7.75 | 0.68 |
| | **dACC** | **Skull** | 38.57 | 256.62 | 2.18 | 0.36 |

| | | | | | | |
|---|---|---|---|---|---|---|
| | | **Soft Tissue** | 37.92 | 477.54 | 7.54 | 0.68 |
| | | **Sphere** | 37.18 | 477.54 | 7.54 | 0.68 |
| | **r-aIns** | **Skull** | 39.12 | 325.69 | 3.51 | 0.46 |
| | | **Soft Tissue** | 38.16 | 460.95 | 7.04 | 0.65 |
| | | **Sphere** | 37.17 | 460.95 | 7.04 | 0.65 |
| **sub-12** | **l-aIns** | **Skull** | 38.42 | 304.47 | 3.06 | 0.43 |
| | | **Soft Tissue** | 37.77 | 375.92 | 4.69 | 0.53 |
| | | **Sphere** | 37.13 | 375.92 | 4.69 | 0.53 |
| | **dACC** | **Skull** | 38.37 | 269.19 | 2.37 | 0.38 |
| | | **Soft Tissue** | 37.79 | 486.25 | 7.83 | 0.69 |
| | | **Sphere** | 37.19 | 486.25 | 7.83 | 0.69 |
| | **r-aIns** | **Skull** | 38.45 | 284.84 | 2.68 | 0.4 |
| | | **Soft Tissue** | 37.76 | 467.49 | 7.24 | 0.66 |
| | | **Sphere** | 37.18 | 467.49 | 7.24 | 0.66 |
| **sub-13** | **l-aIns** | **Skull** | 38.36 | 247.35 | 2.02 | 0.35 |
| | | **Soft Tissue** | 37.83 | 522.02 | 9.01 | 0.74 |
| | | **Sphere** | 37.22 | 522.02 | 9.01 | 0.74 |
| | **dACC** | **Skull** | 39.03 | 327.53 | 3.54 | 0.46 |
| | | **Soft Tissue** | 38.07 | 481.7 | 7.69 | 0.68 |
| | | **Sphere** | 37.19 | 481.7 | 7.69 | 0.68 |
| | **r-aIns** | **Skull** | 39.71 | 405.18 | 5.41 | 0.57 |

| | | | | | | |
|---|---|---|---|---|---|---|
| | | **Soft Tissue** | 38.56 | 464.55 | 7.12 | 0.66 |
| | | **Sphere** | 37.21 | 464.55 | 7.12 | 0.66 |
| **sub-14** | **l-aIns** | **Skull** | 39.52 | 379.79 | 4.75 | 0.54 |
| | | **Soft Tissue** | 38.5 | 504.52 | 8.43 | 0.71 |
| | | **Sphere** | 37.2 | 504.52 | 8.43 | 0.71 |
| | **dACC** | **Skull** | 38.74 | 377.96 | 4.77 | 0.53 |
| | | **Soft Tissue** | 37.93 | 498.46 | 8.24 | 0.7 |
| | | **Sphere** | 37.22 | 498.46 | 8.24 | 0.7 |
| | **r-aIns** | **Skull** | 38.7 | 290.86 | 2.79 | 0.41 |
| | | **Soft Tissue** | 37.97 | 443.15 | 6.51 | 0.63 |
| | | **Sphere** | 37.17 | 443.15 | 6.51 | 0.63 |
| **sub-15** | **l-aIns** | **Skull** | 38.75 | 286.77 | 2.72 | 0.41 |
| | | **Soft Tissue** | 38 | 407.36 | 5.49 | 0.58 |
| | | **Sphere** | 37.09 | 206.21 | 1.45 | 0.29 |
| | **dACC** | **Skull** | 38.78 | 271.51 | 2.43 | 0.38 |
| | | **Soft Tissue** | 37.93 | 383.02 | 4.85 | 0.54 |
| | | **Sphere** | 37.12 | 383.02 | 4.85 | 0.54 |
| | **r-aIns** | **Skull** | 38.77 | 303.42 | 3.06 | 0.43 |
| | | **Soft Tissue** | 37.84 | 423.97 | 5.94 | 0.6 |
| | | **Sphere** | 37.14 | 423.97 | 5.94 | 0.6 |
| **sub-16** | **l-aIns** | **Skull** | 39.22 | 371.1 | 4.56 | 0.52 |

| | | | | | | |
|---|---|---|---|---|---|---|
| | | **Soft Tissue** | 38.36 | 538.41 | 9.61 | 0.76 |
| | | **Sphere** | 37.26 | 538.41 | 9.61 | 0.76 |
| | **dACC** | **Skull** | 39.12 | 417.79 | 5.72 | 0.59 |
| | | **Soft Tissue** | 38.22 | 444.33 | 6.55 | 0.63 |
| | | **Sphere** | 37.19 | 444.33 | 6.55 | 0.63 |
| | **r-aIns** | **Skull** | 38.88 | 308.17 | 3.15 | 0.44 |
| | | **Soft Tissue** | 38.14 | 453.6 | 6.82 | 0.64 |
| | | **Sphere** | 37.19 | 453.6 | 6.82 | 0.64 |
| **sub-17** | **l-aIns** | **Skull** | 38.45 | 277.3 | 2.54 | 0.39 |
| | | **Soft Tissue** | 37.93 | 555.16 | 10.19 | 0.79 |
| | | **Sphere** | 37.24 | 555.16 | 10.19 | 0.79 |
| | **dACC** | **Skull** | 38.3 | 251.97 | 2.05 | 0.36 |
| | | **Soft Tissue** | 37.72 | 477.21 | 7.52 | 0.67 |
| | | **Sphere** | 37.17 | 477.21 | 7.52 | 0.67 |
| | **r-aIns** | **Skull** | 38.27 | 247.99 | 2.03 | 0.35 |
| | | **Soft Tissue** | 37.76 | 484.35 | 7.78 | 0.68 |
| | | **Sphere** | 37.21 | 484.35 | 7.78 | 0.68 |
| **sub-18** | **l-aIns** | **Skull** | 39.5 | 395.32 | 5.09 | 0.56 |
| | | **Soft Tissue** | 38.3 | 340.8 | 3.85 | 0.48 |
| | | **Sphere** | 37.1 | 340.8 | 3.85 | 0.48 |
| | **dACC** | **Skull** | 38.67 | 330.51 | 3.6 | 0.47 |

| | | | | | | |
|---|---|---|---|---|---|---|
| | | **Soft Tissue** | 37.91 | 471.87 | 7.38 | 0.67 |
| | | **Sphere** | 37.2 | 471.87 | 7.38 | 0.67 |
| | **r-aIns** | **Skull** | 40.31 | 463.83 | 7.05 | 0.66 |
| | | **Soft Tissue** | 38.8 | 431.88 | 6.16 | 0.61 |
| | | **Sphere** | 37.16 | 420.77 | 5.87 | 0.6 |
| **sub-19** | **l-aIns** | **Skull** | 40.87 | 431.49 | 6.2 | 0.61 |
| | | **Soft Tissue** | 38.83 | 412.8 | 5.64 | 0.58 |
| | | **Sphere** | 37.15 | 412.8 | 5.64 | 0.58 |
| | **dACC** | **Skull** | 40.29 | 656.15 | 14.29 | 0.93 |
| | | **Soft Tissue** | 39.19 | 487.58 | 8.12 | 0.69 |
| | | **Sphere** | 37.22 | 475.76 | 7.51 | 0.67 |
| | **r-aIns** | **Skull** | 39.22 | 350.57 | 4.07 | 0.5 |
| | | **Soft Tissue** | 38.22 | 525.17 | 9.09 | 0.74 |
| | | **Sphere** | 37.2 | 525.17 | 9.09 | 0.74 |
| **sub-20** | **l-aIns** | **Skull** | 37.35 | 353.48 | 4.12 | 0.5 |
| | | **Soft Tissue** | 37.14 | 442.88 | 6.49 | 0.63 |
| | | **Sphere** | 37.03 | 442.88 | 6.49 | 0.63 |
| | **dACC** | **Skull** | 38.3 | 208.87 | 1.45 | 0.3 |
| | | **Soft Tissue** | 37.66 | 443.29 | 6.49 | 0.63 |
| | | **Sphere** | 37.15 | 443.29 | 6.49 | 0.63 |
| | **r-aIns** | **Skull** | 37.27 | 363.94 | 4.38 | 0.51 |

| | | | | | | |
|---|---|---|---|---|---|---|
| | | **Soft Tissue** | 37.13 | 485.81 | 7.83 | 0.69 |
| | | **Sphere** | 37.03 | 485.81 | 7.83 | 0.69 |
| **sub-21** | **l-aIns** | **Skull** | 39.19 | 333.48 | 3.69 | 0.47 |
| | | **Soft Tissue** | 38.18 | 445.44 | 6.57 | 0.63 |
| | | **Sphere** | 37.16 | 445.44 | 6.57 | 0.63 |
| | **dACC** | **Skull** | 38.5 | 324.87 | 3.4 | 0.46 |
| | | **Soft Tissue** | 37.95 | 506.1 | 8.49 | 0.72 |
| | | **Sphere** | 37.21 | 506.1 | 8.49 | 0.72 |
| | **r-aIns** | **Skull** | 38.91 | 339.77 | 3.82 | 0.48 |
| | | **Soft Tissue** | 38.01 | 446.73 | 6.62 | 0.63 |
| | | **Sphere** | 37.18 | 446.73 | 6.62 | 0.63 |
| **sub-22** | **l-aIns** | **Skull** | 39.18 | 330.06 | 3.6 | 0.47 |
| | | **Soft Tissue** | 38.25 | 491.73 | 8 | 0.7 |
| | | **Sphere** | 37.19 | 491.73 | 8 | 0.7 |
| | **dACC** | **Skull** | 38.47 | 260.68 | 2.22 | 0.37 |
| | | **Soft Tissue** | 37.74 | 409.71 | 5.56 | 0.58 |
| | | **Sphere** | 37.14 | 409.71 | 5.56 | 0.58 |
| | **r-aIns** | **Skull** | 38.87 | 324.06 | 3.46 | 0.46 |
| | | **Soft Tissue** | 38.17 | 530.16 | 9.26 | 0.75 |
| | | **Sphere** | 37.21 | 530.16 | 9.26 | 0.75 |
| **sub-23** | **l-aIns** | **Skull** | 39.62 | 389.63 | 5.04 | 0.55 |

| | | | | | | |
|---|---|---|---|---|---|---|
| | | **Soft Tissue** | 38.47 | 487.64 | 7.88 | 0.69 |
| | | **Sphere** | 37.21 | 487.64 | 7.88 | 0.69 |
| | **dACC** | **Skull** | 38.56 | 296.83 | 2.92 | 0.42 |
| | | **Soft Tissue** | 37.81 | 399.1 | 5.28 | 0.56 |
| | | **Sphere** | 37.15 | 399.1 | 5.28 | 0.56 |
| | **r-aIns** | **Skull** | 39.17 | 332.56 | 3.67 | 0.47 |
| | | **Soft Tissue** | 38.12 | 443.36 | 6.52 | 0.63 |
| | | **Sphere** | 37.18 | 443.36 | 6.52 | 0.63 |
| **sub-24** | **l-aIns** | **Skull** | 39.44 | 369.05 | 4.49 | 0.52 |
| | | **Soft Tissue** | 38.39 | 552.77 | 10.03 | 0.78 |
| | | **Sphere** | 37.25 | 552.77 | 10.03 | 0.78 |
| | **dACC** | **Skull** | 38.2 | 220.79 | 1.62 | 0.31 |
| | | **Soft Tissue** | 37.63 | 476.63 | 7.51 | 0.67 |
| | | **Sphere** | 37.17 | 476.63 | 7.51 | 0.67 |
| | **r-aIns** | **Skull** | 39.07 | 354.26 | 4.13 | 0.5 |
| | | **Soft Tissue** | 38.24 | 481.34 | 7.65 | 0.68 |
| | | **Sphere** | 37.17 | 481.34 | 7.65 | 0.68 |
| **sub-25** | **l-aIns** | **Skull** | 39.17 | 386.45 | 4.94 | 0.55 |
| | | **Soft Tissue** | 38.17 | 430.81 | 6.13 | 0.61 |
| | | **Sphere** | 37.14 | 430.81 | 6.13 | 0.61 |
| | **dACC** | **Skull** | 39.2 | 368.03 | 4.45 | 0.52 |

| | | | | | | |
|---|---|---|---|---|---|---|
| | | **Soft Tissue** | 38.19 | 423.3 | 5.94 | 0.6 |
| | | **Sphere** | 37.18 | 423.3 | 5.94 | 0.6 |
| | **r-aIns** | **Skull** | 39.55 | 381.04 | 4.81 | 0.54 |
| | | **Soft Tissue** | 38.44 | 492.6 | 8.03 | 0.7 |
| | | **Sphere** | 37.2 | 492.6 | 8.03 | 0.7 |
| **sub-26** | **l-aIns** | **Skull** | 40.2 | 401.5 | 5.32 | 0.57 |
| | | **Soft Tissue** | 39 | 407.68 | 5.51 | 0.58 |
| | | **Sphere** | 37.2 | 407.68 | 5.51 | 0.58 |
| | **dACC** | **Skull** | 38.62 | 330.4 | 3.52 | 0.47 |
| | | **Soft Tissue** | 37.93 | 427.66 | 6.06 | 0.6 |
| | | **Sphere** | 37.16 | 427.66 | 6.06 | 0.6 |
| | **r-aIns** | **Skull** | 39.37 | 372.2 | 4.54 | 0.53 |
| | | **Soft Tissue** | 38.37 | 495.01 | 8.08 | 0.7 |
| | | **Sphere** | 37.21 | 495.01 | 8.08 | 0.7 |
| **sub-27** | **l-aIns** | **Skull** | 38.93 | 364.37 | 4.36 | 0.52 |
| | | **Soft Tissue** | 38.13 | 525.36 | 9.13 | 0.74 |
| | | **Sphere** | 37.23 | 525.36 | 9.13 | 0.74 |
| | **dACC** | **Skull** | 38.8 | 313.42 | 3.19 | 0.44 |
| | | **Soft Tissue** | 38.03 | 423.31 | 5.93 | 0.6 |
| | | **Sphere** | 37.15 | 423.31 | 5.93 | 0.6 |
| | **r-aIns** | **Skull** | 39.13 | 387.83 | 4.92 | 0.55 |
| | | **Soft Tissue** | 38.18 | 489.92 | 7.96 | 0.69 |

| | | | | | | |
|---|---|---|---|---|---|---|
| | | **Sphere** | 37.22 | 489.92 | 7.96 | 0.69 |
| **sub-28** | **l-aIns** | **Skull** | 38.69 | 287.72 | 2.73 | 0.41 |
| | | **Soft Tissue** | 37.99 | 520.12 | 8.92 | 0.74 |
| | | **Sphere** | 37.19 | 520.12 | 8.92 | 0.74 |
| | **dACC** | **Skull** | 39.05 | 370.08 | 4.49 | 0.52 |
| | | **Soft Tissue** | 37.98 | 500.3 | 8.28 | 0.71 |
| | | **Sphere** | 37.2 | 500.3 | 8.28 | 0.71 |
| | **r-aIns** | **Skull** | 39.08 | 300.88 | 3 | 0.43 |
| | | **Soft Tissue** | 38.29 | 503.44 | 8.37 | 0.71 |
| | | **Sphere** | 37.19 | 503.44 | 8.37 | 0.71 |
| **sub-29** | **l-aIns** | **Skull** | 38.78 | 277.54 | 2.54 | 0.39 |
| | | **Soft Tissue** | 38.08 | 503.49 | 8.38 | 0.71 |
| | | **Sphere** | 37.21 | 503.49 | 8.38 | 0.71 |
| | **dACC** | **Skull** | 38.46 | 333.03 | 3.57 | 0.47 |
| | | **Soft Tissue** | 37.85 | 464.65 | 7.16 | 0.66 |
| | | **Sphere** | 37.18 | 464.65 | 7.16 | 0.66 |
| | **r-aIns** | **Skull** | 39.98 | 463.05 | 7.02 | 0.65 |
| | | **Soft Tissue** | 38.72 | 458.69 | 6.97 | 0.65 |
| | | **Sphere** | 37.19 | 458.69 | 6.97 | 0.65 |

**Suppl. Table 1** Average temperature, pressure, ISPPA, and MItc per area and per participant in skull, soft tissue, and sphere. The sphere refers to a 10 mm radius spherical mask centred on the default MNI coordinates chosen for each area.

**Supplementary Material 1**

**Absent = Not present**

**Mild = Present but not bothersome**
**Moderate = Tolerable - required some intervention/medication, but did not interfere with day-to-day activities**
**Severe = Intolerable - required contact with a GP or hospital A&E**
Use the middle box to specify whether you think it might be related to the stimulation.
Use the box to the right of each symptom to provide details. e.g. describe what you felt, how long it lasted, medication you took to relieve the symptoms.

| **Since your stimulation, have you had...? symptom to stimulation** | **Intensity of** | **Relationship** | **Please provide details** |
|---|---|---|---|
| a headache | absent | unrelated | |
| neck pain | mild | unlikely | |
| tooth pain | moderate | possible | |
| unusual feelings on your head or scalp | severe | probable | |
| itchiness | | definite | |

changes to your hearing speech problems
vision problems (e.g. double vision) unusual twitching or muscle
movement difficulties in balance
changes in the movement of your
strongest hand
numbness or tingling sensations muscle tightness of the
face or arm unusual feelings, attitudes or emotions
anxiety, worried thoughts or nervousness increased
sleepiness changes to your sleep pattern
difficulty paying attention increased forgetfulness
nausea or sickness to the stomach
dizziness or light-headedness a seizure within the last
24 hours other symptom:

| |
|---|
| other symptom: |
| **Do you have anything else to report?** |

### Supplementary Material 2

### Contraindications

#### 1. Contraindications to Magnetic Resonance Imaging (MRI)

##### 1.1 Intraocular Foreign Bodies (IOFB)

- Individuals with a history of eye trauma involving metal must have an IOFB X-ray prior to MRI.
- MRI is strictly contraindicated unless IOFB has been ruled out via imaging. Applicable to:
- Patients
- Staff
- Students
- Visitors
- Volunteers
- Individuals must not enter the scan room if there are reports of accidents involving metal penetrating the eye and no clearance has been obtained.

##### 1.2 Implants

***General Guidelines***

All implants must be labelled as one of the following:

- **MR SAFE** • **MR CONDITIONAL** • **MR UNSAFE**

**MR CONDITIONAL and MR UNSAFE implants** require an alert on the patient's CRIS record and documented manufacturer information.

Scanning radiographers are responsible for verifying implant safety and modifying scan parameters if necessary.

- If implant safety cannot be confirmed, **do not scan**.
- **All adverse incidents** involving implants and MRI must be reported to:

- (Medicines and Healthcare products Regulatory Agency (**MHRA)** (external)
- **DATIX** (internal)

***Clinical Implants (University Hospitals Plymouth NHS Trust)***

MRI scans must follow specific local procedures (DOPs) including:

- **Intracranial aneurysm clips** (DOP MRI_4) • **Programmable shunts** (DOP MRI_5) • **Active cardiac implants** (DOP MRI_13) • **Hearing implants** (DOP MRI_17) • **Intracranial microsensors** (DOP MRI_21)

**Note:** If the patient has an implant that can be safely scanned on the 0.31T low field strength (extremity) scanner, but not on the 1.5T field scanner, this must clearly be indicated via an ALERT on Clinical Radiology Information System (CRIS).

***Research Participation (University of Plymouth)***

- Presence of any implant is an **exclusion criterion** for healthy volunteer studies.
- Exception: If full implant details are provided by clinical teams and the **MR Safety Expert approves**, scanning may proceed under recommended conditions.

#### 1.3 Pregnancy

***Patients***

- All female patients will be asked if there is **any possibility of pregnancy**.
- The **date of last menstrual period (LMP)** is recorded to prompt further questioning if overdue.
- If pregnancy is possible but denied with confidence by the patient, the scan may proceed.
- MRI during pregnancy must follow **DOP MRI: 7** protocol. ***Staff***
- Pregnant staff should **avoid entering the scan room** during scanning.
- First trimester: Optional exclusion from **MRI environment (inner controlled area)**.
- Non-authorised MRI personnel must not enter the scan room if pregnant.

**Research Scans**

- **Pregnant participants** can only be scanned with **full ethical approval**

### 2. Contraindications to Transcranial Ultrasound Stimulation (TUS)

#### 2.1 Risk Groups and Vulnerable Populations *Pregnancy*

Although magnetic fields (e.g., in repetitive transcranial magnetic stimulation (rTMS)) attenuate quickly with distance and have been safely used in pregnant women (Klírová et al., 2008), no TUS studies have been conducted in this population.

Safety Strategy: Pregnant individuals will be excluded from all TUS studies until peer-reviewed research confirms safety in this group. If inclusion is necessary, it must be justified with specific ethical approval.

***Children and Adolescents***

Despite the safe application of TMS in > 1100 paediatric cases (Frye et al., 2008), the developing brain is physiologically and pharmacologically different from adults.

Safety Strategy: Participants under the age of 18 will be excluded from TUS studies. Inclusion of minors would require specific ethics committee approval and scientific justification.

#### 2.2 Pre-existing Medical Conditions

TUS-induced neural perturbation may be less predictable in participants with existing neurological disorders, posing additional risk.

***Exclusion Criteria (Medical History)***

Participants with the following conditions will be excluded due to heightened seizure risk or unpredictable neuronal reorganization (Rosa et al., 2004):

- Epilepsy or a first-degree relative with idiopathic (non-acquired) epilepsy
- Brain tumour
- Stroke
- Meningitis
- Encephalitis
- Severe head trauma Safety Strategy:

- All participants will be screened for these conditions using a detailed medical questionnaire.

Individuals meeting any of these criteria will not be enrolled.

#### 2.3 Medications and Other Drug Use

The effect of TUS on individuals taking medication remains uncertain, particularly due to the potential interaction with the blood-brain barrier (BBB) or lowered seizure thresholds.

Note: TUS protocols used in our studies do not open the BBB. However, we adopt a cautious approach.

***General Rule***

Participants currently taking any medications will be excluded, including prescription, over-the-counter, or recreational drugs.

Participants must abstain from alcohol for at least 24 hours before the stimulation session.

***High-Risk Drug Categories (based on rTMS seizure risk)*** *Category*

*1 – High Seizure Risk (strong exclusion):*

- Tricyclic antidepressants (e.g., amitriptyline, imipramine)
- Antipsychotics (e.g., clozapine, chlorpromazine)
- Stimulants and illicit substances (e.g., amphetamines, 3,4-methylenedioxymethamphetamine (MDMA), cocaine, ketamine, phencyclidine (PCP), gamma-hydroxybutyrate GHB)

- Certain antivirals and antibiotics (e.g., ganciclovir, ritonavir, foscarnet, theophylline)

*Category 2 – Moderate Seizure Risk (relative exclusion):*

- SSRIs (e.g., fluoxetine, sertraline, paroxetine)
- Anticonvulsants and immunosuppressants (e.g., cyclosporine, lithium)
- Antihistamines, sympathomimetics, anticholinergics
- Antimalarials (e.g., chloroquine, mefloquine) *Category 3 – Withdrawal Risk:*

- Barbiturates
- Benzodiazepines
- Meprobamate
- Chloral hydrate

Safety Strategy:

- Participants taking or recently withdrawing from any of these substances will be excluded.
- Medication status will be assessed via screening forms prior to participation.

#### 2.4 Seizure Risk

***General Considerations***

Although TUS has not been associated with seizure induction, safety measures are informed by the more extensive rTMS literature, where seizures have occurred, especially under high-frequency protocols or in combination with other risk factors (Conca et al., 2000; Rosa et al., 2004; Rossi et al., 2009).

Risk factors from these reports include:

- Use of pro-convulsant drugs (e.g., fluoxetine)
- Sleep deprivation
- History of epilepsy or brain injury

Safety Strategy for Seizure Prevention:

Individuals with seizure disorders or those at increased risk due to medication or medical history will be excluded.

All TUS protocols will strictly adhere to conservative safety parameters, and participants will be monitored throughout stimulation.
